# Munc13 and SNAP25 dependent tethering plays a key role in synaptic vesicle priming

**DOI:** 10.1101/2022.04.10.487799

**Authors:** Christos Papantoniou, Ulrike Laugks, Julia Betzin, Cristina Capitanio, José Javier Ferrero, José Sánchez-Prieto, Susanne Schoch, Nils Brose, Wolfgang Baumeister, Benjamin H. Cooper, Cordelia Imig, Vladan Lučić

**Affiliations:** Department of Molecular Structural Biology, Max Planck Institute of Biochemistry, 82152 Martinsried, Germany; Centre for Structural Systems Biology (CSSB), Heinrich Pette Institut, Leibniz-Institut für Experimentelle Virologie, DESY, 22607 Hamburg, Germany; Department of Neuropathology, University Hospital of Bonn, 53127 Bonn, Germany; Departamento de Bioquímica y Biología Molecular, Facultad de Veterinaria, Universidad Complutense, 28040 Madrid, Spain; Department of Molecular Neurobiology, Max Planck Institute of Multidisciplinary Sciences, City Campus, 37075 Göttingen, Germany; Department of Neuroscience, University of Copenhagen, 2200 Copenhagen, Denmark

## Abstract

Synaptic vesicle tethering, priming, and neurotransmitter release require a coordinated action of multiple protein complexes. While physiological experiments, interaction data, and structural studies of purified systems were essential for our understanding of the function of the individual complexes involved, they cannot combine high structural detail with the unperturbed organization of complexes within cells to resolve how the actions of individual complexes integrate. We employed cryo-electron tomography to simultaneously image multiple presynaptic protein complexes and lipids at molecular resolution in their native composition, conformation and environment. Our results argue that tethers comprising proteins Munc13 and SNAP25 differentially and spatially confine vesicles with single nanometer precision, define vesicle tethering states, and provide molecular mechanisms that guide vesicles towards fusion, which includes molecular priming by conversion to SNARE complex-dependent tethers. These findings present an example of a cellular function performed by an extended molecular assembly comprising multiple, molecularly diverse complexes.

## Introduction

Like most other cellular processes, synaptic vesicle (SV) tethering, priming and fusion require a precise orchestration of molecular pathways that are carried out by multiple macromolecular complexes organized in functional units. The last step of the neurotransmitter release process, SV fusion with the plasma membrane, most often occurs at a specialized region of the presynaptic terminal apposed to the synaptic cleft, termed the active zone (AZ). Prior to fusion, SVs have to be recruited to and held in close proximity to the AZ membrane, involving the tight coupling of the neurotransmitter release machinery, SVs, and voltage-gated calcium channels (Jahn and Fasshauer, 2012; Südhof, 2012, 2013; Rizo, 2018; Brunger et al., 2018).

The molecular mechanisms operating at AZs have long been a focus of nerve cell biology. Here, the Munc13-1 and Munc13-2 members of the Munc13 protein family are necessary for SV priming, i.e. for the formation of a readily releasable pool of SVs, a physiologically defined process that renders individual SVs capable of fusion upon Ca^2+^ influx into the presynaptic terminal (Augustin et al., 1999; Varoqueaux et al., 2002). The RIM protein family has a central role in the organization of the AZ (Schoch et al., 2002; Südhof, 2012). Importantly, RIM1α reverses Munc13-1 homodimerization, thus activating Munc13-1 (Deng et al., 2011). Munc13s and Munc18-1 facilitate the formation of a complex of the three synaptic soluble N-ethylmaleimide-sensitive factor attachment protein receptor (SNARE) fusion proteins (SNAP25, syntaxin-1 and synaptobrevin-2), which is predicted to link SVs with the plasma membrane (Ma et al., 2013). The ultimate fusion of primed SVs is driven by the full assembly of the SNARE complex, a coiled-coil structure comprising four α-helices contributed by synaptic SNAREs (Jahn and Scheller, 2006; Südhof, 2013). Munc13-1 is thought to execute its SNARE-regulating priming function via its MUN-domain, and it is stimulated by Ca^2+^-calmodulin binding to an amphipathic helical motif, by diacylglycerol (DAG) binding to a C_1_ domain, and by Ca^2+^-phospholipid binding via the central C_2_B domain (Rhee et al., 2002; Junge et al., 2004; Shin et al., 2010; Lipstein et al., 2013; Liu et al., 2016; Lipstein et al., 2021). Reconstitution assays showed that efficient liposome fusion requires C_1_, C_2_B, MUN and C_2_C domains of Munc13-1, leading to the notion that Munc13-1 can simultaneously bind SVs and plasma membranes and SNAREs (Rizo, 2018).

Our current understanding of presynaptic AZ function, as briefly outlined above, is based on comprehensive cell biological and functional studies, typically involving the combination of targeted genetic perturbations with morphological, ultrastructural, and electrophysiological analyses. However, the mechanism by which individual steps of the transmitter release process are organized and coordinated in the synapse, the identity of protein machines and their precise mode of operation during individual steps of the process, and the processes that position these machines in the presynaptic compartment for optimal function remain subjects of speculation (Rizo, 2018). One major knowledge gap is how the molecular SV tethering, priming and fusion machinery are organized to drive activity-dependent membrane trafficking and neurotransmitter release. Addressing this issue experimentally in a cellular context is not trivial because it necessitates visualizing both lipids and protein complexes at molecular resolution in situ within the complex environment of the presynaptic AZ.

The vast majority of our insights into synaptic ultrastructure stems from transmission electron microscopy (EM) studies of dehydrated and plastic-embedded synapses. Multiple steps in the preparation of such samples (i.e. chemical fixation, dehydration, heavy metal staining, plastic-embedding, and mechanical sectioning) can introduce structural artifacts that alter or conceal ultrafine molecular arrangements in the synapse. A dense network of filaments interconnecting and tethering SVs was first observed by EM in preparations based on quick freezing and deep-etching, which led to the proposal that these filaments may constrain the movement and cluster SVs (Landis et al., 1988; Hirokawa et al., 1989; Gotow et al., 1991). The widespread implementation of vitrification by high pressure freezing followed by freeze-substitution (HPF/FS) has proven powerful in limiting the extent to which such synaptic artefacts become manifest (Korogod et al., 2015). Ultrastructural analyses of presynaptic vesicle pools in mutant synapses have focused predominantly on measuring distances between vesicles and the plasma membrane, or on the distribution of long, filamentous structures that are detectable with heavy metal-based staining protocols (Siksou et al., 2007, 2009; Imig et al., 2014; Cole et al., 2016; Chakrabarti and Wichmann, 2019). Specifically, deletion of Munc13 or individual SNARE proteins in synapses alters the distribution of vesicles at the AZ and reduces the number of “docked” vesicles (Weimer et al., 2006; Hammarlund et al., 2007; Siksou et al., 2009; Imig et al., 2014), defined as making an apparent physical contact with the AZ plasma membrane in transmission electron micrographs from dehydrated and plastic-embedded samples. Because these preparations cause alterations that preclude molecular interpretation, a major unresolved question in this context remains whether”docked” vesicles are kept in this position through interactions of the SNARE proteins in trans.

In this context, a new methodological approach based on the molecular-level visualization of the presynaptic terminal by cryo-electron tomography (cryo-ET) (Zuber and Lucic, 2019) provided a way forward. This method is uniquely suited for simultaneous, label-free imaging of unstained molecular complexes at single nanometer resolution in situ, i.e. in their native environment (Lucic et al., 2005a; Oikonomou and Jensen, 2017). Indeed, corresponding studies showed, for instance, (i) that pleomorphic, membrane-bound complexes organize SVs by interlinking (via molecular connectors) and tethering them (via molecular tethers) to the presynaptic plasma membrane, (ii) that the distance between SVs and the AZ membrane and the tethering state of the SVs are indicative of SV progression towards fusion (Fernández-Busnadiego et al., 2010, 2013), and (iii) that tethers aid in localizing neurotransmitter release sites near postsynaptic receptors (Martinez-Sanchez et al., 2021). To be precise and avoid the nomenclature ambiguity present in the field, all SVs that are linked to the AZ membrane by molecular bridges (tethers) are called tethered SVs. We refrain from using “docking” because in cryo-ET, SVs are not seen making direct, extended contact with the plasma membrane (Zuber and Lucic, 2019).

Most of these studies were performed on synaptosomes, a well-established model to study the neurotransmitter release process and aspects of the postsynaptic response (Nicholls and Sihra, 1986; Whittaker, 1993; Godino et al., 2007; Dunkley et al., 2008; Martinez-Sanchez et al., 2021), which preserve the macromolecular architecture of neuronal synapses and are excellent substrates for cryo-ET imaging at the molecular level (Schrod et al., 2018; Tao et al., 2018; Zuber and Lucic, 2019; Liu et al., 2019). However, constitutive knockout mice lacking key components of the presynaptic neurotransmitter release machinery typically die at birth, i.e. at a developmental stage were hardly any synapses are formed in the forebrain, so that the purification of high-quality synaptosomes with acceptable yield is essentially impossible.

In the present study, we solved this problem by implementing experimental strategies for cryo-ET analysis of synaptosomal preparations obtained from organotypic slice cultures from embryonic mouse brains. We therefore performed structural molecular imaging of priming-deficient synapses by cryo-ET to resolve the mechanism for organizing AZ-proximal SVs, determine the vesicle tethering functions of Munc13s, SNAP25, and RIMs and directly identify some of the tethers. Based on our findings, we propose a new model of the molecular mechanisms mediating the SV progression towards fusion, where RIMs, Munc13s, and SNAREs are required in subsequent steps that lead from SV tethering, via priming, to fusion.

## Results

### Munc13 and SNAP25 do not affect the overall distribution of synaptic vesicles

Deletion of Munc13 and SNAP25 proteins in neurons leads to complete abolition and severe impairment of release, respectively, and a loss of fusion-competent vesicles (Schoch et al., 2001; Varoqueaux et al., 2002; Washbourne et al., 2002), as well as a redistribution of synaptic vesicles in the vicinity of the AZ plasma membrane (Hammarlund et al., 2007; Siksou et al., 2009; Imig et al., 2014).

Because Munc13 and SNAP25 knockout mice die at birth, we proceeded to establish an experimental workflow that would allow us to perform cryo-ET on synapses from cultured material. Specifically, we prepared hippocampal organotypic slice cultures at embryonic day 18 (E18) and cultured the tissue for 3-5 weeks before synaptosome preparation. During the culture time, slices could recover from the dissection trauma and develop mature synapses – also in material derived from Munc13- and SNARE-deficient mouse mutants (Imig et al., 2014; Sigler et al., 2017). We detected and imaged synaptosomes directly in the crude synaptosomal fractions from organotypic slices. These could not be visually distinguished form gradient centrifugation-based synaptosomes, with the added advantage that avoiding the gradient centrifugation step shortened the preparation time as compared to our previous studies (Fernández-Busnadiego et al., 2013).

Cryo–electron tomograms of synaptosomes showed smooth and continuous membranes, spherical SVs at a distance to the plasma membrane, a well-defined synaptic cleft, and no signs of cytoplasmic aggregation (Figure 1A), as expected from vitrified samples and in accordance with previous studies (Supplementary video 1) (Zuber and Lucic, 2019). Each presynaptic terminal contained many SVs, a mitochondrion and an apposed postsynaptic terminal containing a postsynaptic density characteristic of excitatory synapses.

**Figure 1:**
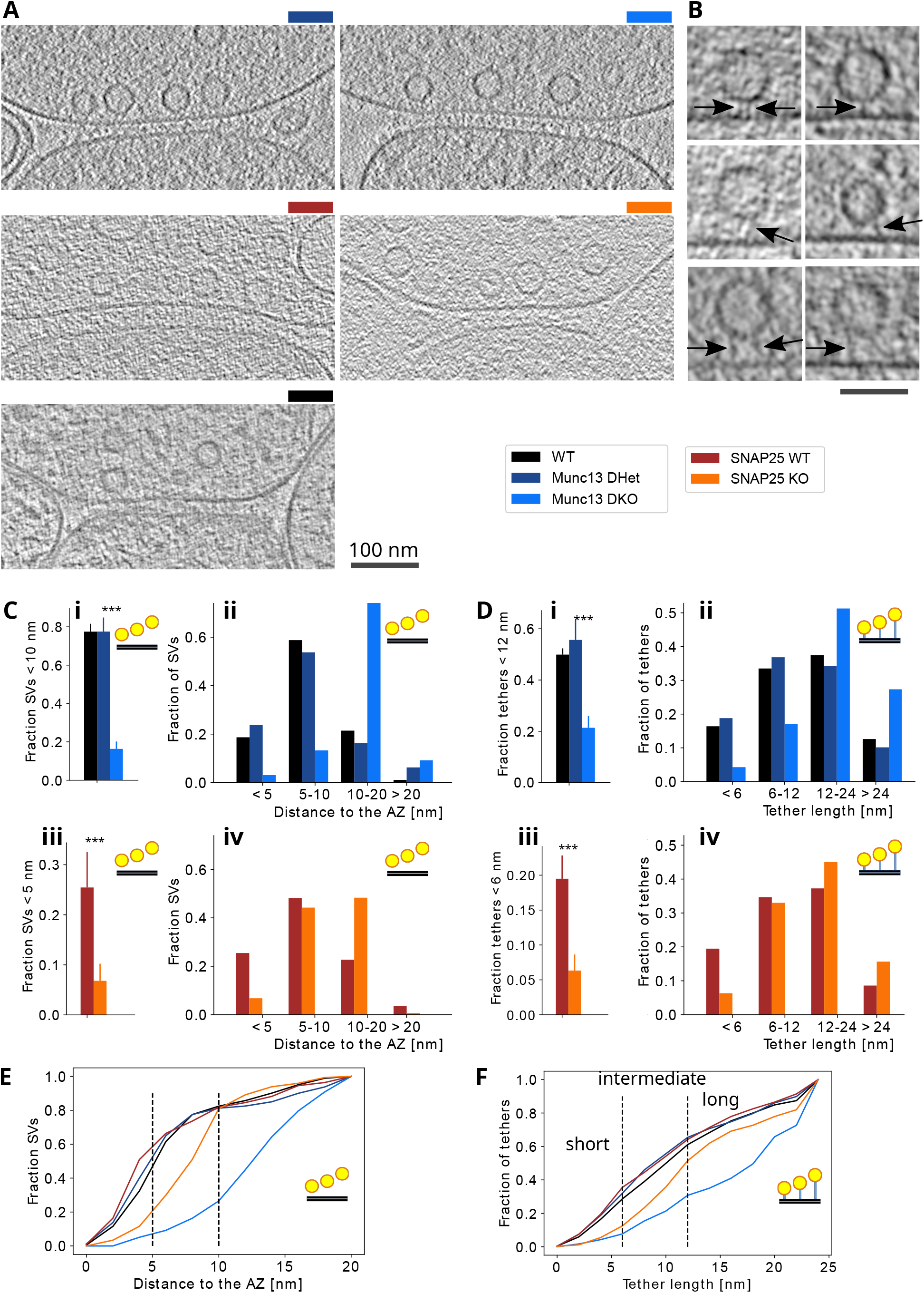
Influence of Munc13 and SNAP25 on the proximal SV location and tether length. (A) Cryo-ET slices of synapses from different conditions, as indicated by the legend. Scale bar 100 nm. (B) Magnified cryo-ET slices showing examples of proximal SVs and tethers: SVs located <5 nm / 5-10 nm / further than 10 nm to the AZ membrane and short / intermediate / long tethers are shown on the top / middle / bottom panels The images are from WT (top and middle) and Munc13 DKO synapses (bottom). Arrows point to tethers. Scale bar 50 nm. (C) Fraction of proximal SVs <10 nm (above) and 5 nm (below) to the AZ membrane and histograms of the proximal SV distances where bins represent SV distance classes. (D) Fraction of tethers shorter than 12 nm (above) and 6 nm (below) to the AZ membrane, and histograms of tether lengths where bins represent short, intermediate and long tethers. (E) Cumulative distribution function of the SV distances, where the distances used to classify the proximal SV distances are shown by black, dashed lines. The data shown here are the same as the data shown in histograms in C. (F) Cumulative distribution function of tether lengths, where the lengths used to classify tethers are shown by black, dashed lines. The data shown here are the same as the data shown in histograms in D.

We compared synapses from mice lacking both dominant Munc13 isoforms (Munc13-1 and Munc13-2) (Munc13 DKO) (Supplementary video 2) and Munc13 double heterozygous (DHet) littermate controls. As this breeding strategy did not enable us to obtain littermate controls that were wildtype for both Munc13 alleles, we prepared and processed organotypic slices from C57BL6/N mice in parallel (WT). In this case, cultures were generated from animals on the day of birth (postnatal day 0), and not at E18. We further imaged hippocampal organotypic slice synaptosomes from mice lacking SNAP25 (SNAP25 KO) (Washbourne et al., 2002) and their littermate controls (SNAP25 WT).

In all conditions imaged, SV concentration profiles showed an accumulation of vesicles in the proximal zone, that is within a distance of 45 nm from the AZ membrane (termed proximal SVs), a decreased concentration in the intermediate zone (45-75 nm to the AZ membrane) and again an increased concentration in the distal zone (75-250 nm to the AZ zone) (Figure S1Ai). We previously observed that a perturbation of this pattern indicates a massively disturbed SV organization and neurotransmitter release (Fernández-Busnadiego et al., 2010, 2013). Furthermore, the surface concentration of proximal SVs of synapses lacking Munc13 and SNAP25 was very similar to their controls (Munc13 DHet and SNAP25 WT, respectively) (Figure S1Aii). The higher concentration observed in the WT condition in comparison to other control conditions was likely caused by the different genetic background or the different age of the animal on the day of culturing (E18 vs P0).

In sum, genetic ablation of Munc13 and SNAP25 neither changed the overall organization of SVs, nor their concentration at the proximal zone.

### Munc13 and SNAP25 organize AZ-proximal synaptic vesicles

We then investigated whether Munc13 and SNAP25 influence the SV organization within the proximal zone. The distance between proximal SVs and the AZ membrane was 14.3 ± 0.4 nm (mean ± sem) in Munc13 DKO synapses, significantly larger than in Munc13 DHet (8.4 ± 0.6 nm, t-test p<0.001) and WT (8.3 ± 0.3 nm) synapses (Figure S1Bi). Probability distributions (normalized histograms) of the SV distances showed peaks located around 16 nm for Munc13 DKO and around 6 nm for Munc13 DHet synapses, indicating that there are at least two AZ distance-dependent SV states (Figure S1Biii).

While the histograms based on 2 nm bin size clearly displayed the separation of the SV distance peaks (Figure S1Biii), the same data plotted at 5 nm bin size show more clearly that the effect of Munc13 removal is most prominent at 0-10 nm from the AZ (Figure 1Cii). The cumulative distribution function of the distances further revealed that the distance threshold between these two peaks was at *∼*10 nm (Figure 1E). To confirm that the distance of 10 nm provides a valid separation of the two states, we calculated the fraction of proximal SVs in the two states. We found that only 15% of proximal SVs in Munc13 DKO synapses were located <10 nm to the AZ membrane, while this was the case for about 80% (that is 4.7 times more SVs) in Munc13 DHet and WT synapses, and that this difference was highly significant (p<0.001, χ^2^ test) (Figure 1Ci). In line with this finding, the fraction of proximal SVs closer than 5 nm was 7.6 times larger in Munc13 DHet than in Munc13 DKO synapses (p<0.001, χ2 test) (Figure S1Bii). Together, our data indicate that Munc13-deletion leads to a shift in the distribution of SVs of approximately 10 nm away from the AZ.

We found that the removal of SNAP25 increases the mean proximal SV distance to the AZ (from 8.2 ± 0.5 nm in SNAP25 WT to 9.8 ± 0.3 nm in SNAP25 KO, t-test p=0.002) (Figure S1Ci). As for the Munc13 conditions, histograms based on 2 nm bins showed distinct peaks (Figure S1Ciii). However, the most pronounced difference was found at approximately 5 nm, as judged by the cumulative SV distance distributions and 5 nm bin histograms (Figure 1E, Civ). Consistent with these results, the fraction of SVs located <5 nm to the AZ membrane was significantly reduced in the absence of SNAP25 (3.7 times smaller, from 25% in SNAP25 WT to 7% in SNAP25 KO synapses, p<0.001 χ^2^ test) (Figure 1Ciii), while the fraction of SVs <10 nm to the AZ membrane was 1.4 times smaller in SNAP25 KO synapses (p<0.001, χ^2^ test) (Figure S1Cii).

Here, we defined the characteristic SNAP25 and Munc13 SV distances (5 and 10 nm, respectively) using the optimal histogram peak separations values (Figures 1E, and S1Biii, Ciii). Nevertheless, there is an inherent uncertainty in these values because the fraction of SV closer to AZ calculated for similar values (e.g. 4 and 9 nm) was also significantly different (p<0.001 χ^2^ test for both SNAP25 KO and Munc13 KO synapses, respectively; data not shown).

Taken together, we found that the removal of Munc13 and SNAP25 shifted the distribution of SV distances towards larger distances (Figures 1Cii, Civ, E), in a qualitative agreement with the previous HPF/FS investigation (Imig et al., 2014). In addition, both the spatial extent and the magnitude of the SV depletion in the proximal zone caused by Munc13 deletion were greater than those after SNAP25 removal.

### Munc13 and SNAP25 control the SV tether types

Next, we tested whether Munc13 and SNAP25 affect tether morphology.

Visual inspection confirmed our previous observations that SV tethers and connectors were the main structural elements organizing SVs (Figure 3A) (Fernández-Busnadiego et al., 2010; Zuber and Lucic, 2019). SV tethers were computationally detected in an automated, unbiased manner using hierarchical connectivity segmentation and their length was measured by taking tether curvature into account, as previously described (Lucic et al., 2016). We found that it was significantly increased in Munc13 DKO (20.9 ± 1.2 nm, mean ± sem) compared to Munc13 DHet (13.1 ± 0.5 nm) and SNAP25 KO (15.7 ± 0.5 nm) compared to SNAP25 WT (12.7 ± 0.4 nm) synapses (p<0.001 in both cases, t-test) (Figure S1Di, Ei).

The cumulative distribution of tether lengths showed a clear separation at approximately 6 and 12 nm (Figure 1F), which is 20% larger than the previously defined SV distance limits (5, 10 and 20 nm). Consequently, we classified tethers by their length into <6 nm (short tethers), 6-12 nm (intermediate) and 12-24 nm (long) groups. Using this classification, we found that the fraction of all tethers that are shorter than 12 nm was significantly decreased in Munc13 DKO synapses, as was the fraction of all tethers shorter than 6 nm in SNAP25 KO synapses (p<0.001 in both cases, t-test) (Figure 1Di, Diii), which can be also seen on tether length histograms (Figures 1Dii, Div, S1Diii, Eiii). We also obtained very significant results when tether lengths little different from 6 and 12 nm were used to define tether classes (p<0.001 t-test for both SNAP25 KO at 5 nm and Munc13 DKO at 10 nm).

In sum, our SV tether analysis showed that deletion of Munc13 or SNAP25 strongly and differentially increases tether length, that Munc13 deletion caused stronger effects, and that these changes parallel alterations in AZ-proximal SV localization. Therefore, we defined three Munc13 and SNAP25-dependent SV tethering and localization states. Namely, SVs localized <5 nm to the AZ membrane and characterized by short tethers belong to the SNAP25-dependent state, SVs localized >10 nm to the AZ membrane and long tethers are Munc13-independent, and the SVs localized 5-10 nm to the AZ membrane with intermediate tethers belong to the intermediate state.

### The lack of all major RIM family proteins severely impairs SV organization

Deletion of RIM proteins, core components of the presynaptic AZ, results in a severe impairment of release and a strong reduction in the number of SVs in the vicinity of the AZ zone membrane, as assessed by EM of chemically fixed samples (Kaeser et al., 2011). We examined the role of RIMs (Schoch et al., 2002) in SV organization at the AZ using cryo-ET of synapses lacking all four multidomain RIM isoforms, 1α, 1β, 2α and 2β. Because mice deficient for all RIM1 and RIM2 variants die after birth (Kaeser et al., 2011), we prepared synaptosomes from cultured primary neurons of double-floxed mice (RIM1/2 Flox/Flox) and compared synapses from neurons expressing active (RIM cDKO) or inactive (RIM Ctrl) Cre-recombinase.

The mean SV distribution profiles in RIM Ctrl synapses showed the expected features, i.e. an accumulation of SVs in the proximal zone and the lower SV concentration in the intermediate zone, while both of these features were lacking in RIM cDKO samples (Figure S2A). Inspection of SV profiles in individual synapses showed that none of the five RIM cDKO synapses had these features, contrary to all four RIM Ctrl synapses assessed.

Furthermore, the distribution of the proximal SV distances to the AZ membrane was shifted towards higher distances in RIM cDKO synapses as compared to Ctrl synaptosomes (p=0.0064, χ2 test) (Figure S2B). The fraction of SV <10 nm to the AZ membrane was significantly smaller in RIM cDKO tomograms (p=0.0036, t-test). The surface concentration of proximal SVs at the AZ membrane was 2.4 times smaller in RIM cDKO synapses, but this difference did not reach significance because of the high variance (p=0.18, t-test).

Our data indicate that the overall synaptic distribution of SVs and their concentration in the proximal zone is severely disturbed in synapses lacking all RIM1 and RIM2 isoforms, even more so than what we had observed previously in RIM1α KO synapses (Fernández-Busnadiego et al., 2013).

### PDBu supports the SNAP25-dependent state

Phorbol esters, including 4-β-phorbol-12,13-dibutyrate (PDBu), are analogs of the endogenous second messenger DAG and known to bind and activate Munc13 and other C1 domain proteins, leading to a potentiation of neurotransmitter release in a Munc13-dependent manner (Newton, 1995; Iwasaki et al., 2000; Rhee et al., 2002). To elucidate structural correlates of this potentiation, we analyzed cryo-ET images of untreated and PDBu-treated neocortical synaptosomes from adult wildtype mice. Furthermore, we also examined synapses treated with PDBu together with either PKC inhibitor Ro31-8220, or Calphostin C, an inhibitor of DAG/PDBu binding to C_1_ domains of Munc13 and PKC.

Analyses of neocortical synaptosomes in the absence of stimulation confirmed that PDBu increases glutamate release. This effect was prevented by Calphostin C, but not by specifically inhibiting PKCs using Ro31-8220 (Figure S3A), as shown previously (Martín et al., 2011). Furthermore, glutamate release from PDBu-treated synaptosomes stimulated by 5 mM KCl was increased upon the addition of extracellular Ca^2+^(Figure S3B).

We then tested whether application of PDBu influences SV tethering as assessed by cryo-ET. We subjected synaptosomes to an extracellular solution that did not contain Ca^2+^ to prevent spontaneous vesicle fusion after PDBu potentiation. We found that neither PDBu alone nor PDBu in combination with Calphostin C or Ro31-8220 affected the overall SV distribution or the surface concentration of proximal SVs (Figure S4A). However, the proximal SV distance to the AZ membrane was significantly decreased by PDBu (p=0.037, t-test) (Figure S4B) and this decrease was caused by the significantly higher fraction of proximal SVs of PDBu-treated synapses located <5 nm to the AZ membrane (p=0.0017, χ^2^-test) (Figure 2Bi, Bii). The lack of significant changes upon combined Calphostin C +PDBu treatment indicates that the PDBu-induced SV translocation we observed is mediated by Munc13 C_1_ domain activation.

**Figure 2:**
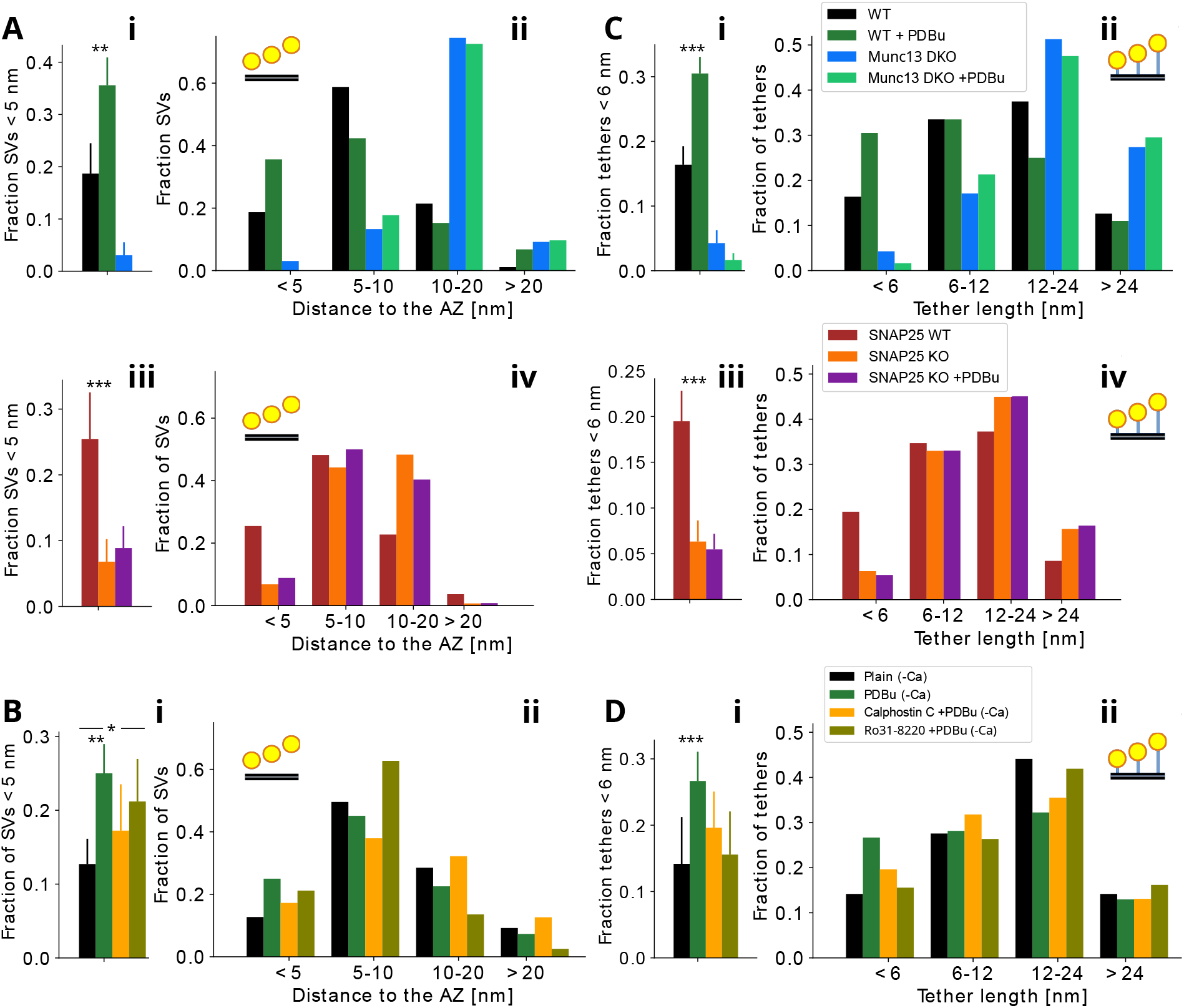
Influence of PDBu on the proximal SV localization and tether length. (A) Fraction of proximal SVs located <5 nm to the AZ membrane (left) and histograms of SV localizations for Munc13 and SNAP25 conditions (right). (B) The same as A, but for pharmacologically treated neocortical synaptosomes. (C) Fraction of short tethers (<6 nm) (left) and tether length histograms (right) for Munc13 and SNAP25 conditions. (D) The same as C, but for pharmacologically treated neocortical synaptosomes.

Furthermore, the proximal SV distance to the AZ membrane was strongly decreased by combined Ro31-8220 +PDBu as compared to control values (p<0.001, t-test), even more so than by PDBu alone (Figure S4B). This change was evident for SV distances up to 10 nm from the AZ membrane, as can be seen from the histogram of proximal SV distances (Figure 2Bii) and from the data showing that the fraction of Ro31-8220 +PDBu proximal SV is significantly increased for SVs <5 nm (Figure 2Bi), and very significantly increased for SVs <10 nm to the AZ membrane (p=0.040 and p<0.001 respectively, χ^2^-test) (Figure S4B). Together, these observations indicate that PKC primarily affects SVs within 5-10 nm from the AZ membrane.

Tether length was significantly decreased by PDBu (p=0.013, t-test) (Figure S4C). This decrease was most prominent for short tethers (p<0.001, χ^2^-test) (Figure 2Di, Dii) and less so but still significant for tethers up to 12 nm in length (p=0.003, χ^2^-test) (Figure S4C). Neither Calphostin C +PDBu nor Ro31-8220 +PDBu resulted in significant differences.

In sum, we found that PDBu reduces the proximal SV distance to the AZ and the tether length in a way that supports the SNAP25-dependent SV tethering state, thus having an effect opposite of the one caused by deletion of SNAP25.

### PDBu-mediated regulation of SV localization and tether length is Munc13- and SNAP25-dependent

To clarify the molecular mechanism of the PDBu effects on SV localization and tethers, we analyzed Munc13 and SNAP25 deficient synapses after PDBu treatment. As expected, the application of PDBu on WT synaptosomes caused a significant increase in fractions of proximal SVs <5 nm to the AZ membrane (p=0.007, t-test) and short tethers (p=0.005, t-test) (Figure 2Ai, Aii, Ci, Cii). These data are very similar to those presented in the previous section (Figure 2B, D) although different extracellular Ca^2+^ concentrations were used (i.e. 1.2 mM Ca^2+^ here vs. nominally Ca^2+^ in previous section).

In contrast, we found that neither the localization of SVs within the proximal zone, nor their tether lengths were changed in PDBu-treated Munc13 DKO and SNAP25 KO synapses (Figures 2A, 2C, S4D, S4E). There were no changes for any of the SV distance and tether length bins.

Therefore, these data confirm the observation that PDBu supports the SNAP25-dependent SV tethering state, and that this effect requires both Munc13 and SNAP25.

### Munc13, SNAP25 and PDBu increase SV tethering

We previously showed that the increase in both tether length and the number of tethers bound to an SV are indicative of the SV progression towards fusion (Fernández-Busnadiego et al., 2010). Therefore, we tested whether the genetic and pharmacological treatments we employed so far influence this relationship and quantitatively determine the interaction between tethers and SVs.

We found that the number of tethers per proximal SV was reduced by 66% in Munc13 DKO synapses (p<0.001, Kruskal-Wallis test, as compared to Munc13 DHet) (Figure 3Ai). The fraction of proximal SVs that are tethered and the fraction of proximal SVs that are multiply tethered (three or more tethers) were very significantly decreased in Munc13 DKO synapses (p<0.001 in both cases, χ^2^-test) (Figure S5Bi, Bii). A large fraction of proximal SVs did not harbor any tethers. Accordingly, the distribution of the number of tethers per SV is strongly shifted towards 0-2 tethers per SV (Figure S5Ai).

**Figure 3:**
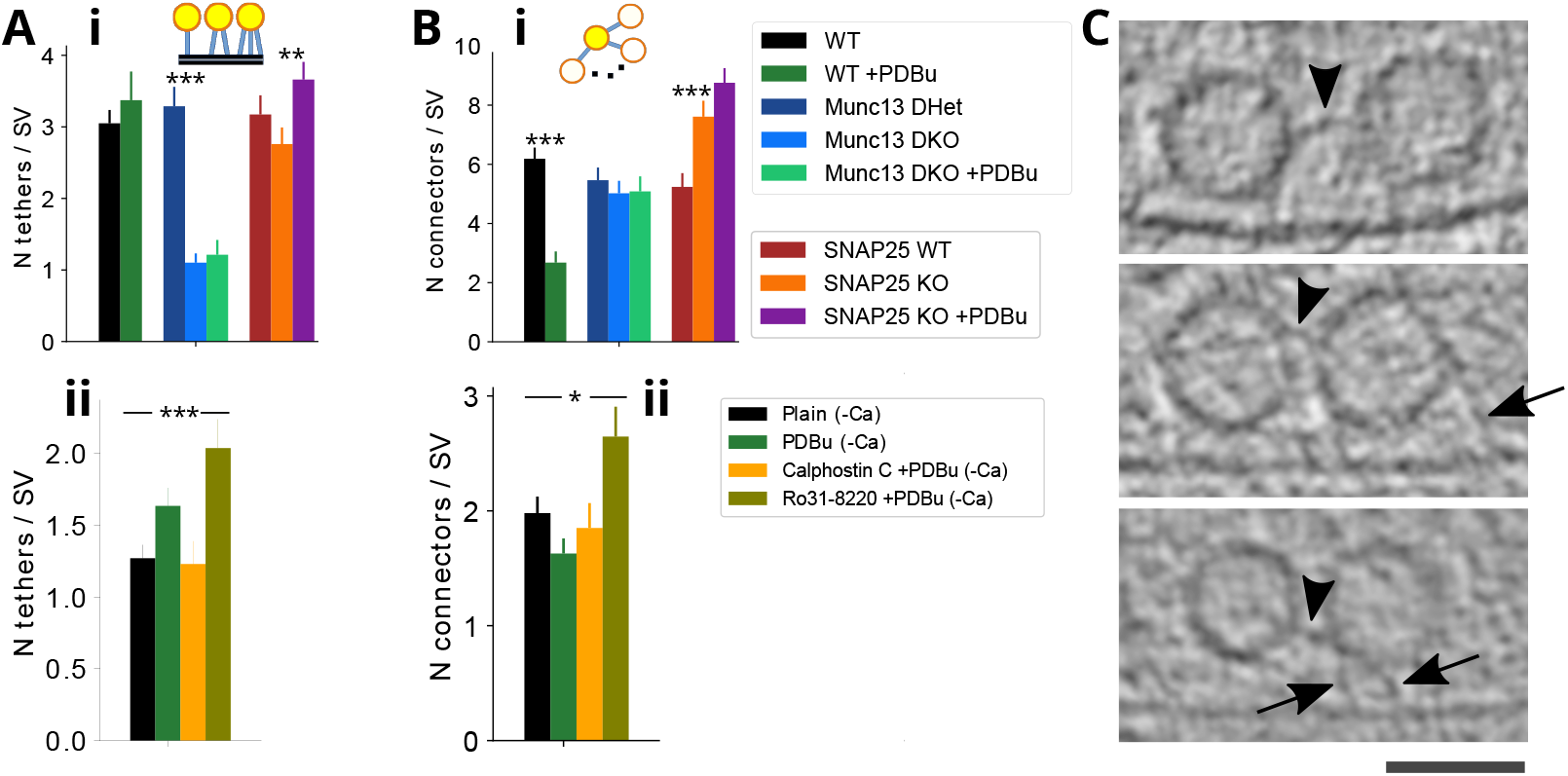
SV tethers and connectors. (A) Number of tethers per proximal SV for Munc13 and SNAP25 conditions (above) and pharmacologically treated neocortical synaptosomes (below). (B) Number of connectors per proximal SVs for the same conditions as in A. (C) Cryo-ET slices showing examples of SV connectors. Arrows point to tethers and arrowheads to connectors. Scale bar 50 nm.

In SNAP25 KO synapses, the mean number of tethers per proximal SV (Figure 3Ai), the fraction of proximal SVs that were tethered (Figure S5Bi), and the fraction of proximal SVs that exhibited multiple tethers (Figure S5Bii) were not altered in comparison to SNAP25 WT synapses. However SNAP25 KO synapses exhibited an increase in the fraction of SVs with only few (1-2) tethers (Figure S5Aii) and a significant decrease in the number of tethers per tethered SV (p=0.015, Kruskal-Wallis test) (Figure S5Biii).

Coapplication of PDBu and the PKC inhibitor Ro31-8220 to WT synapses increased the number of tethers per proximal SV, the fraction of tethered proximal SVs, and the fraction of multiply tethered proximal SVs (p<0.001 Kruskal-Wallis test, p=0.021 χ^2^-test and p=0.027 χ^2^-test, respectively, all in comparison to WT) (Figures 3Aii, S5Bi, S5Bii, respectively). PDBu alone, however, did not significantly increase the number of tethers per proximal SV either in the presence or absence of extracellular Ca^2+^ (Figure 3Ai, Aii). In line with our observation that PDBu treatment of synapses lacking Munc13 did not rescue the changes in SV distribution and tether length (Figures 2A, C, S4D, E), we found that the decrease in the mean number of tethers per proximal SV (Figure 3Ai), the fraction of tethered SVs, and the fraction of SVs that exhibited multiple tethers seen in absence of Munc13 (Figure S5Bi, Bii) were not reverted by PDBu. In contrast, PDBu treatment of synapses lacking SNAP25 very significantly increased the number of tethers per SV (p=0.003, Kruskal-Wallis test) (Figure 3Ai). This increase was mostly caused by an increased fraction of proximal SVs with multiple tethers (p=0.003, χ^2^-test) (Figure S5Bii) and it reverted the peak at 1-2 tethers per SV observed in the distribution of the number of tethers per SV in SNAP25 KO tomograms (Figure S5Aii). This finding is surprising, given that PDBu treatment of SNAP25 KO synaptosomes did not rescue the shift of SVs to larger distances from the AZ membrane (Figures 2Aiii, Aiv, S4D) nor the tether length (Figures 2Ciii, Civ, S4E).

Together, these results indicate that SNAP25 increases the number of tethers per tethered SV, while Munc13, and to a lesser extent PKC inhibition, increase both the fraction of tethered SVs and the number of tethers. The effect of Munc13 had a wider range and was stronger than that of SNAP25. Surprisingly, in the absence of SNAP25, PDBu treatment caused an increase in the number of tethers per SV without affecting tether length, thus decoupling the inverse relationship between tether length and the number of tethers per SV that we observed before (Fernández-Busnadiego et al., 2010).

### SNAP25, PDBu and PKC decrease the proximal SV connectivity

We previously found that tethering alterations observed upon certain genetic manipulations are correlated with modifications of SV connectors (bridges interlinking SVs) (Fernández-Busnadiego et al., 2013; Vargas et al., 2017). Therefore, we investigated whether proximal SV connectors contribute to the SV tethering states defined above.

The connectors were visualized and automatically detected using the same procedure we used for tethers (Figure 3C). The number of connectors per proximal SV was significantly decreased by PDBu treatment of WT synapses in the presence of extracellular Ca^2+^ and significantly increased in SNAP25 KO samples (p<0.001 in both cases, Kruskal-Wallis test) (Figure 3 Bi). However, neither the removal of Munc13, nor the application of PDBu on Munc13 DKO or SNAP25 KO synapses caused significant changes in the number of connectors per proximal SV (Figure 3 Bi). Furthermore Ro31-8220 +PDBu induced a significant increase in the number of connectors per proximal SV (p=0.034, Kruskal-Wallis test), but PDBu alone did not, both in the absence of extracellular Ca^2+^, (Figure 3Bii).

We proceeded to determine the locus of the observed SV connectivity changes. We found that the PDBu- and Ro31-8220-induced alterations were highly significant for SVs located at 5-10 nm from the AZ membrane (p<0.001 in both cases, Kruskal-Wallis test) (Figure S5Ci, Ciii), whereas the increase in SNAP25 KO synapses affected SVs within 5-10 and 10-20 nm (p=0.010 and p=0.009, respectively, Kruskal-Wallis test) (Figure S5Cii). Furthermore, despite the lack of a PDBu effect on the number of connectors per SV for all proximal SVs taken together in the absence of extracellular Ca^2+^(Figure 3B), there was a significant decrease for SVs within the 10-20 nm region upon PDBu and Calphostin C +PDBu treatments (p=0.002 and p=0.003, respectively, Kruskal-Wallis test) (Figure S5Ciii).

In sum, connector properties followed a pattern distinct from the one observed for tethers. Namely, our data indicate that PDBu, SNAP25 and PKC, but not Munc13, act to reduce the number of connectors per SV. All three factors showed a significant connectivity decrease for SVs located 5-10 nm to the AZ, indicating that changes in SV connectivity preferentially affect SVs in the intermediate state.

### Munc13 and SNAP25 deletion affect SV size

We found that the radius of proximal SVs was significantly increased in Munc13 DKO and SNAP25 KO synapses (p<0.001 in both cases, t-test) (Figure S5Di), in agreement with previous observations in Munc13- and SNAP25-deficient synapses from HPF/FS organotypic slices (Imig et al., 2014). In addition, PDBu decreased the radius of SVs in WT synaptosomes in the presence and absence of extracellular Ca^2+^, and in SNAP25 KO synapses (p<0.001, p = 0.003 and p=0.019, respectively, t-test), but not in Munc13 DKO synapses (Figure S5Di, Diii). This phenomenon did not exclusively affect proximal SVs, but persisted for all analyzed SVs (within 250 nm to the AZ membrane) (Figure S5Dii).

Therefore, among the conditions tested, smaller SV size correlated with treatments that increase neurotransmitter release and larger SVs with conditions known to disturb neurotransmitter release. The precise mechanisms by which SV size and functional neurotransmitter release properties might be linked are unclear (Imig et al., 2014).

### Synapses from intact neurons

We pursued two strategies to image synapses of intact neurons of dissociated rat E17-21 hippocampal cultures that were grown on electron microscopy grids and vitrified at DIV 21. First, we searched for synapses in regions distant from cell bodies that were sufficiently thin to allow direct cryo-ET imaging (roughly up to 500 nm). These thin regions contained predominantly non-synaptic axonal boutons, as observed previously (Schrod et al., 2018).

Second, we employed focused ion beam milling at cryo-conditions (cryo-FIB) (Marko et al., 2007; Rigort et al., 2012; Schaffer et al., 2017), applying wedge-shaped and lamella geometries to reduce sample thickness in the vicinity of cell bodies. We pursued both lamella and wedge cryo-FIB milling geometries (Figure S6A). However, these procedures suffer from synapse-specific difficulties such as long culturing times on EM grids, the requirement to optimize culture density to maintain stability of milled regions while insuring proper vitrification, and FIB-induced material redeposition (Fernandez et al., 2016). All these factors together severely limited the number of synapses that could be located and imaged by cryo-ET, resulting in a total of five tomograms.

We checked that neurons were properly vitrified and had a healthy appearance, as done for synaptosomes (Figure S6B; Supplementary video 3). In addition, we made sure that pre- and postsynaptic membranes formed an extended synaptic cleft of approximately 25 nm in width, as observed before in vitrified synapses (Lucic et al., 2005b; Tao et al., 2018). This was necessary to properly identify synapses because neurons in culture often have non-synaptic axonal boutons that contain many SVs and have their plasma membrane closely apposed to a different process at a distance that is too small to be a synaptic cleft (Krueger et al., 2003; Schrod et al., 2018).

Importantly, the SV organization, the lack of extended membrane contacts between SVs and the AZ, connectors and tethers in neuronal cultures were very similar to those observed by cryo-ET in synaptosomes (Figure S6B) (Lucic et al., 2007; Schrod et al., 2018; Tao et al., 2018; Liu et al., 2020). Quantitative analysis of the overall SV distribution relative to the AZ membrane showed a well-defined peak in the proximal and a minimum in the intermediate zone (Figure S6C). Furthermore, the distributions of proximal SV distances to the AZ membrane and tether lengths, were consistent with those of synaptosomes (Figures 1C, D).

### Fitting atomic models into tethers

Biochemical and structural investigations showed that Munc13 and the SNARE complex can bind two lipid membranes simultaneously (Jahn and Südhof, 1999; Liu et al., 2016). We therefore examined whether the tethers we detected in our tomograms contain these proteins, by rigid body fitting of the relevant currently available atomic models into the tethers.

First, we used an atomic model from a crystal structure of the primed SNARE complex, which is thought to link primed vesicles to the AZ membrane [(Zhou et al., 2017) PDB id: 5w5d] (Figure 4A). This model comprises one SNARE motif each of VAMP2 and Syntaxin 1a, and two SNARE motifs of SNAP25, together with a helical fragment of Complexin 1 and the C2B domain of Synaptotagmin 1. The four SNARE motifs form a parallel α-helix bundle with their C-terminal ends positioned close to their vesicle (VAMP2) and plasma membrane-associated domains (Syntaxin 1a and SNAP25). This model necessitates a short distance between the SNARE complex-bound SV and the plasma membrane because the C-terminal ends of all four SNARE motifs are very close to their respective membrane anchors.

**Figure 4:**
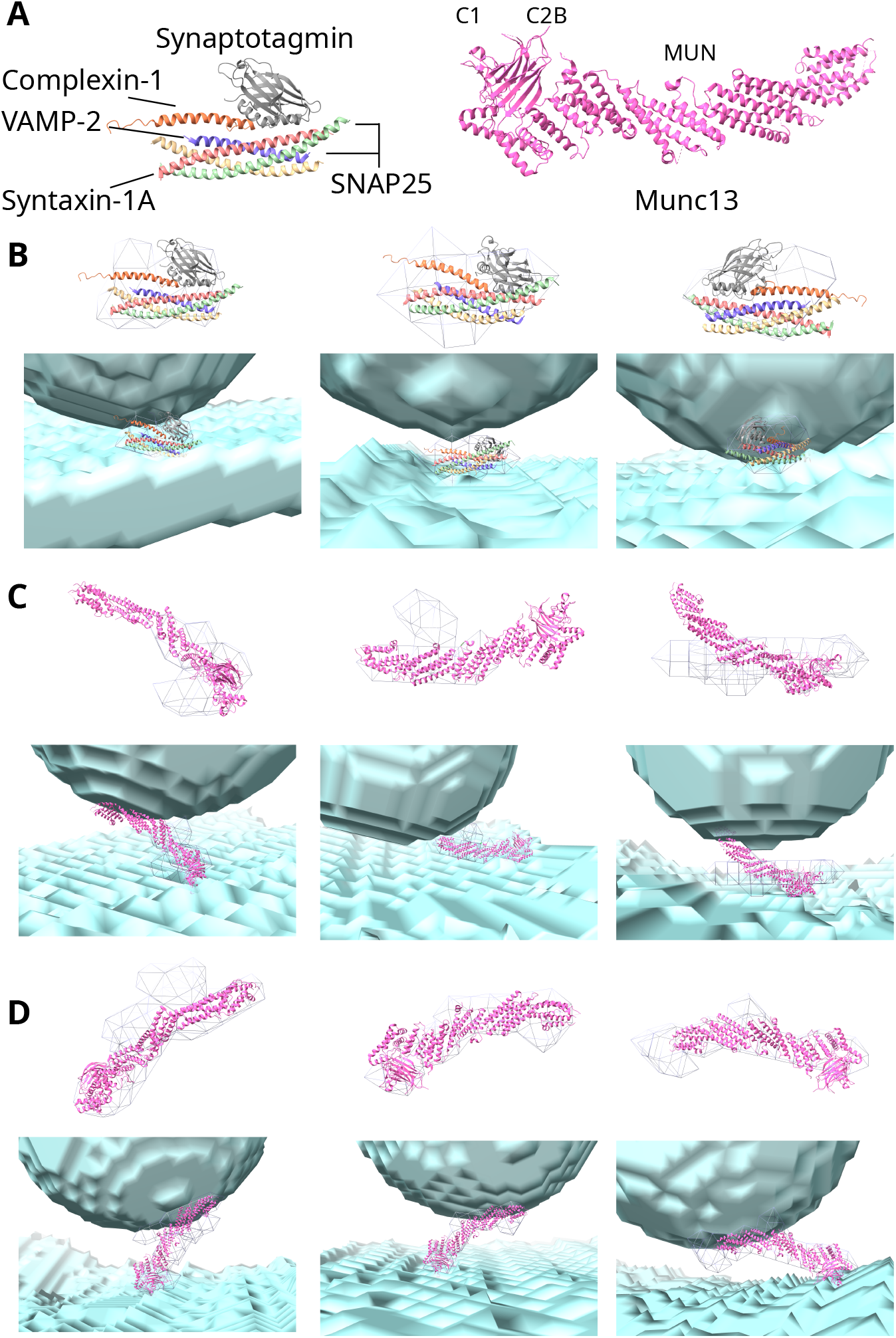
Fitting atomic models to tethers. (A) Atomic models of the primed SNARE complex (PDB id: 5w5d; left) and Munc13 C_1_, C_2_B and MUN domains (PDB id: 5ue8; right). (B) Examples of the primed SNARE complex fitting in short tethers (C) Examples of intermediate tethers where the Munc13 C_1_C_2_BMUN model cannot fit. (D) Examples of tethers where the Munc13 C_1_C_2_BMUN model can fit. (B)-(D) Left column panels show examples from the WT, central column B from SNAP25 WT, central columns C and D from SNAP25 KO and the right column from intact neurons. Grey wire meshes show the segmented tether surface, while the SVs and plasma membrane are shown as semi-transparent cyan-blue volumes. In each panel, the same fit is shown without (above) and with the SVs and plasma membrane (below).

Considering tethers in both segmented and greyscale density forms, we found that this model fits well into many short tethers in synaptosomes and intact neurons (Figure 4B). These tethers properly accounted for the distance between the SV and plasma membranes and contained a lateral density that was adequately placed and sufficiently large to account for the SNARE helices. Other short tethers were large enough to accommodate the primed SNARE complex, but the distance between the SVs and plasma membranes was too long, or they contained an extra density that might correspond to additional protein(s).

The second model we used is the largest currently available atomic model of a Munc13 fragment, comprising C_1_, C_2_B and MUN domains, and covering 55% of the entire Munc13 sequence [(Xu et al., 2017) PDB id: 5ue8]. Because Munc13 has a rod-shaped MUN domain that is flanked by the lipid-binding C_2_B and C_2_C domains (Figure 4A), and the MUN domain is homologous to tethering factors in other systems Munc13 was hypothesized to form tethers (Sztul and Lupashin, 2009; Rizo, 2018; Quade et al., 2019).

A fit was deemed acceptable if the region from the C_2_B domain to the C-terminal end of MUN domain could be placed within the segmented tether. We found that this Munc13 model could fit into many intermediate and long tethers. This might be surprising, considering that the distance between the vesicle and the plasma membrane binding regions of Munc13 is at least 13.0 nm, as determined by measuring the shortest distance between the C_2_B domain and the C-terminal end of the MUN domain of the Munc13 atomic model. However, these tethers contained extended density along the plasma and/or vesicle membranes that made fitting the Munc13 C_1_C_2_BMUN model possible (Figure 4D). Nevertheless, we found that 19 of 49 intermediate tethers in SNAP25 KO synapses were too small to accommodate the Munc13 C_1_C_2_BMUN model (Figure 4C). Furthermore, the model could not fit 18 out of 61 tethers in WT and Munc13 DHet synapses and 16 out of 31 tethers in intact neurons. The fact that the Munc13 C_1_C_2_BMUN model only covers 55% of the Munc13 sequence and does not include binding partners increases the significance of this finding.

In sum, short tethers were sufficiently large to accommodate the primed SNARE complex, and the Munc13 C_1_C_2_BMUN domain fit into many intermediate tethers. However, a significant number of intermediate tethers could not accommodate any of these models. Because these tethers were observed in WT, Munc13 DHet, SNAP25 KO, and in intact neurons, they are most likely stable and of physiological relevance, and at least some of them do not contain SNAP25.

## Discussion

### Methodological considerations

Over the past three decades we have achieved a very detailed understanding of the molecular, morphological, and functional aspects of the neurotransmitter release process and the underlying SV cycle. This progress is mainly owed to genetically modified model organisms lacking defined protein components of the SV cycle. However, gaining detailed insight into the complex molecular machinery that operate the SV trafficking steps preceding fusion, characterizing their precise composition, localization, and interrelation in synapses in vivo, and pinpointing the precise roles of the key molecular players that conduct separate SV trafficking tasks has remained exceedingly difficult, which led to many controversies in the field. Arguably the biggest methodological problem in this context has been that only EM approaches provide the necessary resolution to visualize the distribution of SVs at individual AZs with nanometer precision. Most such approaches require post-fixation dehydration of samples, heavy-metal staining of membrane and cytomatrix components, and embedding in plastic resin for ultramicrotomy. This allows the in-depth characterization of the SV organization relative to the AZ under different experimental conditions and the indirect immunodetection of proteins, but artefacts introduced by the individual processing steps cannot be avoided - so that proteins cannot be seen directly with appropriate stringency.

Cryo-ET, as employed in the present study provides the most promising way forward in this regard. It permits the visualization of synaptic protein complexes in the vitrified frozen hydrated state and without the use of heavy metals to enhance membrane contrast. Most importantly, it enables the visualization of SVs and the AZ plasma membrane bound complexes, thus allowing insights into the native molecular organization of synaptic proteins in minimally perturbed synapses at a single nanometer scale.

Our observations that Munc13 and SNAP25 are needed for SV localization <10 nm and <5 nm to the AZ membrane, respectively, qualitatively agree with the SV distribution results of the same mutants obtained by ET of high pressure frozen / freeze-substituted (HPF/FS) organotypic slices (Siksou et al., 2009; Imig et al., 2014). However, the corresponding distances were shorter in the HPF/FS study. For example, SVs making direct contact with the AZ membrane in HPF/FS samples correspond to SVs located up to 5 nm but not touching the AZ membrane, while SVs of Munc13 DKO synapses accumulated at 8-10 nm to the AZ membrane in HPF/FS and further than 10 nm in our data. This discrepancy is likely due to a single nanometer scale rearrangements caused by dehydration of HPF/FS samples (Henrich et al., 2009; Bleck et al., 2010) or the heavy metal staining. Also, the increase of SV size upon Munc13 and SNAP25 deletion observed in organotypic slices prepared by HPF/FS (Imig et al., 2014) agree with our cryo-ET results from synaptosomes.

Regions of vitrified dissociated neuronal cultures where most synapses are located are mostly too thick for cryo-ET imaging (Schrod et al., 2018), thus necessitating sample thinning by cryo-FIB milling. We observed that the structural features in synapses of intact neurons were consistent with results from synaptosomes, and that membrane contacts between SVs and the AZ were extremely rare in synaptosomes and synapses from intact neurons, in agreement with previous cryo-ET assessments (Lucic et al., 2007; Fernández-Busnadiego et al., 2010; Schrod et al., 2018; Tao et al., 2018; Liu et al., 2020).

Together, the correspondence between our cryo-ET results from synaptosomes with the HPF/FS results from organotypic slices, and the agreement between cryo-ET date from synaptosomes and intact synapses provide orthogonal validations of our synaptosome model to study molecular architecture of synapses.

Here we combined the use of genetic approaches to study synapses [i.e. knockout mice for key regulators of AZ organization (RIMs), SV priming (Munc13s), and fusion (SNAP25)] with cryo-ET of synaptosomes.

While RIM cDKO abolishes expression of all relevant RIMs (Kaeser et al., 2011) and Munc13 DKO abrogates both evoked and spontaneous release (Augustin et al., 1999; Varoqueaux et al., 2002), compensatory effects cannot be excluded in SNAP25 KO synapses because SNAP23 supports asynchronous release and a low level of spontaneous release remains despite severe release impairment (Washbourne et al., 2002; Delgado-Martínez et al., 2007).

### Potential molecular identity and function of SV tethers

We visualized complexes involved in SV priming and neurotransmitter release at a single nanometer resolution in their native composition, conformation and environment, within entire presynaptic terminals. Precise localization, analysis of structural alterations caused by genetic and pharmacological perturbations, and molecular fitting allowed us to identify some of the SV-interacting presynaptic complexes and led us to propose a SV tethering and priming model that combines structural and molecular information.

In line with our previous work (Fernández-Busnadiego et al., 2010, 2013), we found that tether lengths exhibited a wide, molecular conditions–dependent range, arguing that tethers of different molecular composition coexist at the synapse. Furthermore, changes in tether length induced by genetic and pharmacological manipulations paralleled those of SV distance to the AZ membrane. Because under different conditions SVs assumed markedly different, non-uniform spatial distribution, despite their thermally-induced diffusion that acts to make the distribution more uniform, and because we did not observe other structures that could influence this distribution, our data strongly argue that tethers control the SV distance to the AZ.

Fitting atomic models of known molecular candidates for SV tethers, when constrained by the known orientation and location of interacting partners, allowed us to distinguish between the models, clearly identify unsuccessful fits, and also to identify those where a model could fit into a tether with or without additional proteins.

We show here that RIM isoforms other than RIM1α also contribute to localization of SVs in the proximal zone. This indicates that the partial compensation observed earlier in a subset of synapses lacking RIM1α was due to 1β, 2α and/or 2β isoforms (Fernández-Busnadiego et al., 2013). Our data strongly argue that Munc13s (Munc13-1 and Munc13-2) and SNAP25 act to reduce the proximal SV distance to the AZ membrane and tether length with a precision of a few nanometers. Quantitative assessment of this data allowed us to distinguish a Munc13-independent SV tethering state and two Munc13-dependent states, namely the SNAP25-dependent and the intermediate state (Figure 5). Our observation that the alterations of the SV distribution, tether length and the number of tethers per SV caused by the deletion of Munc13 were more pronounced than those of SNAP25 argues that the tethering function of Munc13 is upstream from SNAP25. Therefore, we conclude that regarding SV progression towards fusion, the Munc13-independent state is the most upstream, followed by the intermediate and the SNAP25-dependent state.

**Figure 5:**
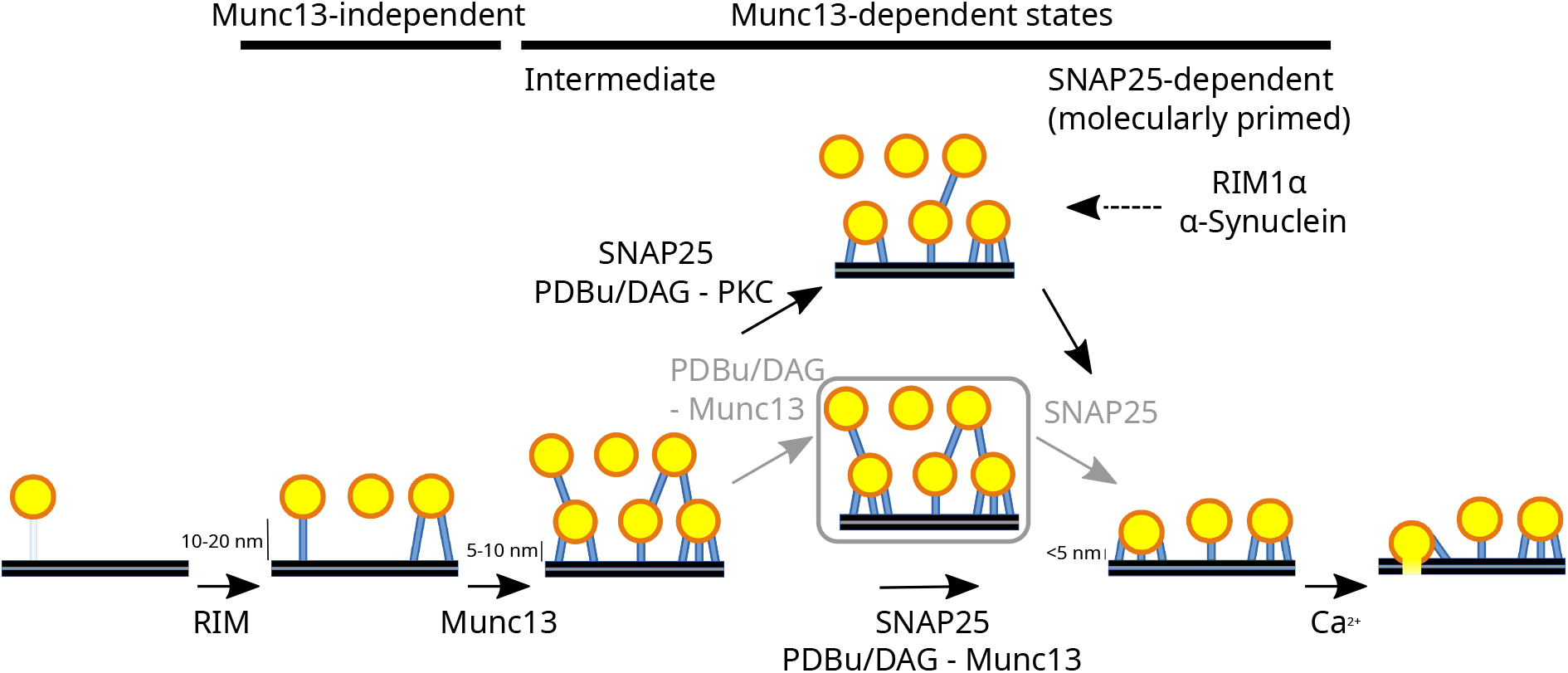
Structural and molecular SV tethering and priming scheme. All SV distances to the AZ membrane and the molecular assignments result from this study, except the influence of RIM1α and α-Synuclein on SV connectors, which was reported earlier (Fernández-Busnadiego et al., 2013; Vargas et al., 2017) and is shown here to provide a complete model. The grey box and grey arrows indicate possible separate contributions of PDBu and SNAP25 to the transition from the intermediate to the SNAP25-dependent state. The connectors-mediated process is shown above. The model is not drawn to scale to emphasize that SVs are located at different distances to the AZ membrane.

These conclusions agree with the known function of Munc13 in facilitating the SNARE complex assembly and with the currently accepted intuitive notion that the SV progression towards fusion proceeds by reducing the SV distance to the AZ membrane (Jahn and Fasshauer, 2012; Rizo, 2018), showing the predictive power of our approach.

The shape and orientation of the majority of intermediate tethers agreed with the atomic model of the Munc13 C_1_C_2_BMUN fragment, previously proposed to bridge SV and plasma membranes (Xu et al., 2017; Quade et al., 2019). Furthermore, given that it was earlier argued that neuronal SNAP25 paralogs are unlikely to significantly compensate for a loss of SNAP25 (Delgado-Martínez et al., 2007; Imig et al., 2014), it is likely that most of the intermediate state SVs are SNAP25-independent. Also considering that the intermediate tethers required Munc13, our data suggest that the majority of intermediate tethers contain Munc13.

The SNAP25-dependent state likely corresponds to the molecularly primed state because it requires Munc13 and SNAP25, proteins necessary for functional priming (Augustin et al., 1999; Varoqueaux et al., 2002; Washbourne et al., 2002). We showed that short tethers strongly depend on the presence of SNAP25, an obligatory component of the SNARE complex, and that their shape and orientation are consistent with the atomic model of the primed SNARE complex (Zhou et al., 2017). This argues that short tethers likely comprise the SNARE complex, in agreement with the fundamental role the SNARE complex plays in membrane fusion (Südhof, 2013; Brunger et al., 2018; Rizo, 2018). This finding does not preclude the presence of partially zippered SNARE complexes, because these are likely morphologically similar to the primed SNARE complex and thus hardly distinguishable in our tomograms.

Furthermore, our data showed that in the presence of Munc13 and SNAP25, DAG/PDBu promotes the transition to the SNAP25-dependent state. In the absence of Munc13, PDBu did not alter the SV localization or tethering, arguing that PDBu directly affects Munc13, as shown before (Rhee et al., 2002). However, in synapses lacking SNAP25, PDBu increased the number of tethers without promoting the transition to the SNAP25-dependent state. This may indicate that SV priming proceeds by a DAG/PDBu-Munc13 dependent formation of intermediate tethers, followed by the SNAP25-dependent formation of short tethers (Figure 5).

Based on our structural data, we propose a sequential SV tethering and priming model comprising the following steps (Figure 5). (i) RIM family members are responsible for the accumulation of SVs in the proximal zone (within 45 nm to the AZ membrane) in a Munc13-independent manner. (ii) Munc13-dependent intermediate tethers are necessary to localize SVs <10 nm to the AZ membrane to the intermediate SV tethering state. (iii) SNAP25-dependent short tethers are needed to localize SVs <5 nm to the AZ membrane into the SNAP25-dependent (molecularly primed) state. The transition to this state is facilitated by DAG/PDBu and requires both the upstream function of Munc13 and SNAP25. (iv) Ca^2+^influx may lead to vesicle fusion.

Together, our data suggest that SV tethers comprising Munc13 and the SNARE complex differentially confine SVs with a single nanometer precision. These results define sequential SV tethering states that precede neurotransmitter release, and provide a molecular mechanism that precisely localizes SVs and explains the observed non-uniform spatial distribution of SVs.

### Possible implications for current models of presynaptic function

While we could clearly distinguish three SV tethering states, the complete picture might be more complex. Specifically, we detected intermediate tethers that were too small to be reconciled with the Munc13 C_1_C_2_BMUN model, and where the distance between their SV and plasma membrane lipid binding regions was too large for the primed SNARE complex model. Because at least some of these tethers did not contain SNAP25, we can speculate that full-length native Munc13 can adopt a more compact structure than that of Munc13 C_1_C_2_BMUN model. While Munc13 showed some flexibility in purified and reconstituted systems (Xu et al., 2017; Gipson et al., 2017; Quade et al., 2019; Grushin et al., 2022), this flexibility is not sufficient to explain the Munc13-incompatible intermediate tethers. Alternatively, these tethers could be formed by a membrane-bridging complex of a size between that of SNARE complex and Munc13 that does not contain SNAP25 or Munc13, but may contain a larger SNAP25 paralogue, or other proteins, as proposed earlier (Jahn and Fasshauer, 2012; Chen et al., 2021),

It has previously been proposed that distinct sub-pools within the overall pool of functionally primed SVs exist (Neher, 2015; Neher and Brose, 2018), which exhibit differences in their probability to fuse in response to an action potential. In this context, DAG/PDBu increases the fraction of high-release probability SVs (also referred to as “superprimed” SVs) relative to the low-release probability primed SVs at the calyx of Held synapses (Taschenberger et al., 2016). Based on our data, we hypothesize that at least some SVs in the SNAP25-dependent SV tethering state correspond to the superprimed functional state. Among the molecularly primed SVs, a transition from low-to a high-release probability state may be achieved by the addition of new SNARE complex-dependent short tethers that may help to pull SVs and AZ membranes together. This is consistent with our earlier hypothesis that a higher number of tethers facilitates SV progression towards fusion (Fernández-Busnadiego et al., 2010; Zuber and Lucic, 2019) and the proposal that small changes in the number of SNARE complexes can regulate priming and release (Arancillo et al., 2013). While the intermediate state SVs are not molecularly equipped to fuse in response to a single action potential, they may rapidly transition into the fusion-competent SNAP25-dependent (molecularly primed) state during synaptic activity - likely in a calcium-dependent and dynamic manner (Neher and Brose, 2018; Chang et al., 2018; Kusick et al., 2020; Maus et al., 2020; Radecke et al., 2022).

Furthermore, we observed that PDBu, SNAP25 and PKC, but not Munc13, decrease the connectivity between proximal SVs. This effect was the most prominent for SVs located 5-10 nm to the AZ membrane, arguing that it is relevant for the transition from the intermediate to the SNAP25-dependent state. The effects of PDBu were reported to be mediated by Munc13 and PKC-dependent pathways (Iwasaki et al., 2000; Rhee et al., 2002; Basu et al., 2007; Wierda et al., 2007; Martín et al., 2010; Genc et al., 2014; Xu et al., 2017; Wang et al., 2021). Therefore, our data suggests that two SNAP25-dependent molecular processes are involved in this transition, while the PDBu-Munc13 process is mediated by tethers, the PDBu-PKC process is mediated by connectors (Figure 5).

Interestingly, Rab3, a SV-associated protein, RIM and Munc13 were early on implicated in functionally superpriming SVs. We previously detected an increased connectivity in synapses lacking RIM1α and in synapses that overexpressed human α-synuclein on a synuclein null background (Schlüter et al., 2006; Fernández-Busnadiego et al., 2013; Vargas et al., 2017). This suggests that RIMs, Rab3, and possibly synucleins, may be involved in the connectors-mediated process. In addition, RIMs were shown to be important in presynaptic plasticity, while PDBu-induced and other forms of plasticity were associated with Rab3 superpriming (Kaeser and Südhof, 2005; Taschenberger et al., 2016). Therefore, we propose that the tether-mediated priming process (Munc13- and SNAP25-dependent) is modulated by the connector-mediated process (SNAP25-dependent and Munc13-independent) and that the latter may be relevant for presynaptic plasticity, thus extending our earlier hypothesis (Zuber and Lucic, 2019).

The relation between SNAP25 and PKC in the connectors-mediated process is unclear. The possibility that PDBu-induced PKC phosphorylation of SNAP25 potentiates synaptic release was recently contested (Shimazaki et al., 1996; Genoud et al., 1999; Ruiter et al., 2020). Alternatively, PKC and SNAP25 could act independently from each other to decrease connectivity. In this scenario, SNAP25 could be associated with SVs in a way that blocks or competes with the formation of connectors. Considering that SNAP25 is also needed to form SNARE complexes, this would require a high concentration of SNAP25 molecules in the close vicinity of a SV undergoing the transition to the SNAP25-dependent state. Such a mechanism could provide a partial explanation for the very high copy number and a wide, SV-associated distribution of SNAP25 within the presynapse (Wilhelm et al., 2014).

Additionally, the opposite roles of SNAP25, increasing the number of tethers and decreasing the number of connectors, present an inverted picture in comparison to synucleins, previously shown to negatively affect tethers and support connectors (Vargas et al., 2017). This may suggest that a concerted, differential regulation of tethers and connectors represents a specific pattern that influences the SV dynamics in the proximal zone.

Finally, we previously described trans-synaptic subcolumns, tripartite protein assemblies that contain tethers and provide a structural link between SVs and postsynaptic receptors, and hypothesized that they ensure efficient synaptic transmission by colocalizing SVs and postsynaptic receptors across the synaptic cleft (Martinez-Sanchez et al., 2021). Furthermore, we showed that subcolumns cluster to form large, non-uniform molecular assemblies, which may be closely related to synaptic nano-columns that were previously described by super-resolution fluorescence imaging (Tang et al., 2016; Chen et al., 2018; Martinez-Sanchez et al., 2021). Our current findings add another functionally relevant layer to the trans-synaptic sub-columns and raise the possibility that connectors interlink subcolumns to form structuraly defined synaptic nano-columns, which organize synaptic transmission. Because these assemblies are intermixed with other complexes and cannot be isolated, they have to be investigated at a single nanometer resolution in situ, thus making our cryo-ET–based approach uniquely suited to achieve this objective. Therefore, our results extend the notion of molecular complexes serving as basic functional modules (Hartwell et al., 1999), and present an example of a cellular process carried out by a concerted action of multiple spatially separated and molecularly diverse complexes comprising a large, non-periodic, protein assembly.

## Materials and Methods

### Munc13- and SNAP25-deficient mouse breeding

Munc13-deficient mice were bred with permission of the Niedersächsisches Landesamt für Verbraucherschutz und Lebensmittelsicherheit (LAVES; 33.19.42502-04-15/1817). All mice were kept according to the European Union Directive 63/2010/EU and ETS 123. They were housed in individually ventilated cages (type II superlong, 435 cm2 floor area; TECHNIPLAST) under a 12 h light/dark cycle at 21 ± 1QC with food and water *ad libitum*, and the health status of the animals was checked regularly by animal care technicians and a veterinarian. Hippocampal organotypic slice cultures were prepared from C57 B6/N wild-type animals on postnatal day (P) 0, and from Munc13- and SNAP25-deficient and littermate control animals at embryonic day 18 (E18) due to perinatally lethal phenotypes (Varoqueaux et al., 2002; Washbourne et al., 2002). E18 mice lacking Munc13-1 and Munc13-2 (Munc13 -/- -/-, denoted in text Munc13 DKO) (Augustin et al., 1999; Varoqueaux et al., 2002) and control littermates were generated by crossing Munc13-1+/-;Munc13-2+/- with Munc13-1+/-;Munc13-2-/- or Munc13-1+/-;Munc13-2+/-mice. Cultures from control littermates with the genotypes Munc13-1+/-Munc13-2+/- (Munc13 DHet) and from P0 control C57 B6N wildtype animals (M13 WT) were prepared on the same day. E18 mice lacking SNAP25 (SNAP25 -/-, denoted in the text SNAP25 KO) were generated by cross-breeding SNAP25+/-mice. Control littermates were wildtype for the SNAP25 allele (SNAP25 WT). We used and pooled data generated from female and male mice since loss of Munc13 or SNAP25 affects synaptic transmission equally in neurons from both genders. That said, we cannot exclude the possibility that subtle changes in the organization of vesicle pools may exist between male and female neurons in wildtype cultures. Analyses were performed 3-5 weeks after culturing. On the day of experiment, slices from animals of the same genotype from cultures prepared on the same day were pooled to get sufficient material for synaptosome preparation.

### Organotypic slice culture preparation from Munc13- and SNAP25-deficient mice

Hippocampal organotypic slice cultures were prepared from Munc13- and SNAP25-deficient and the corresponding controls mice using the interface method (Stoppini et al., 1991) as described previously (Imig et al., 2014; Maus et al., 2020). Pregnant females at E18 were anaesthetized and decapitated and embryos were removed by hysterectomy. Pups were decapitated and hippocampi were dissected in dissection medium (97 ml Hank’s balanced salt solution, 2.5 ml 20% glucose, 1 ml 100 mM kynurenic acid, pH 7.4). Three hundred _μ_m-thick hippocampal slices with the entorhinal cortex attached were prepared using a McIlwain tissue chopper. Slices were then transferred onto sterile Millipore membrane confetti pieces on top of 6-well membrane inserts in 1.2 ml of culture medium (22.44 ml ddH_2_O, 25 ml 2xMEM, 25 ml BME, 1 ml GlutaMAX, 1.56 ml 40% Glucose, 25 ml horse serum). Residual dissection medium was removed using a P200 pipette. A maximum of 4 hippocampal slices were cultured per membrane insert at 37°C and 5% CO_2_. Slice culture medium was changed 24 hours after preparation and then 2-3 times per week.

### Synaptosomal preparation from Munc13- and SNAP25-deficient slice cultures

Munc13- and SNAP25-deficient and the corresponding control synaptosomes were prepared from DIV 28-30 hippocampal organotypic slices. The incubation buffer was removed and the slices were briefly rinsed with Tyrode’s buffer (120 mM NaCl, 3 mM KCl, 1.25 mM MgCl_2_, 1.25 mM CaCl_2_, 0.5 mM NaH_2_PO_4_, 25 mM Hepes, 30 mM D-glucose, pH 7.4) at room temperature. They were scooped with a brush and dropped in a glass tube containing the homogenization buffer (HB; 0.32 M sucrose and one tablet Complete mini EDTA-free protease inhibitor cocktail (Roche; 10 ml, pH 7.4)) at 4°C (50 _μ_l of buffer for each slice). Slices were homogenized in a teflon glass homogenizer applying one stroke at 100rpm and 7 strokes at 700rpm. The homogenate was centrifuged at 2000 g for 2 min (twice), the pooled supernatants were centrifuged for 12 min at 9500 g and the pellet (P2) was resuspended in Ca^2+^ free Hepes-buffered medium (HBM; 140 mM NaCl, 5 mM KCl, 5 mM NaHCO_3_, 1.2 mM NaH_2_PO_4_-H_2_O, 1 mM MgCl_2_-6H_2_O, 10 mM Glucose, 10 mM Hepes, pH 7.4), yielding crude synaptosomal fraction (all at 4°C). The protein concentration was measured using the Bradford Assay (Bio-Rad) on an Amersham Biosciences Ultrospec 3100 Pro Spectrophotometer (GE Healthcare) and diluted to 0.3 – 0.5 mg/ml. The fraction was centrifuged at 10 000 g for 10 min and the pellet was stored on ice.

One hour before vitrification, the pellet containing the synaptosomes was thoroughly and carefully resuspended with warm HBM + 1.2 mM CaCl_2_ and was incubated at 37°C. Some synaptosomes (as indicated in the Results) were treated with 1 _μ_M Phorbol-12,13-dibutyrate (PDBu) in DMSO (final concentration) for 15 min.

### Neocortical synaptosomal preparation

Cerebrocortical synaptosomes were extracted from 6–8 week old male Wistar rats as described previously (Dunkley et al., 1988; Godino et al., 2007; Fernández-Busnadiego et al., 2013) in accordance with the procedures accepted by the Max Planck Institute for Biochemistry. In brief, animals anesthetized with chloroform were decapitated, and the cortex was extracted and homogenized in homogenization buffer (HB; 0.32 M sucrose, 20 mM DTT, and one tablet of Complete mini EDTA-free protease inhibitor cocktail (Roche; 10 ml, pH 7.4)) with up to seven strokes in a Teflon glass motor driven homogenizer at 700 rpm. The homogenate was centrifuged at 2000 g for 2 min (twice), the pooled supernatants were centrifuged for 12 min at 9500 g and the pellet (P2) was resuspended in Ca^2+^ free HBM (all at 4°C), yielding crude synaptosomal fraction. The suspension was loaded onto a three-step Percoll (GE Healthcare) gradient (3%, 10% and 23%, all in 0.32 M sucrose solution), the maximum suspension volume was 1ml per gradient. The gradients were centrifuged in a Beckmann ultracentrifuge at 25000g at 4°C for 6 minutes and the material accumulated at the 10/23% interface was retrieved with a Pasteur pipette and diluted in 50ml of HBM. Percoll was removed by an additional washing step with HBM by centrifugation for 10 min at 22000 g. As we realized that healthy synaptosomes from the crude synaptosomal fraction could be detected and imaged in the same way as the ones prepared by Percoll gradient centrifugation, we decided to omit this step for synaptosomes intended for cryo-ET. Percoll gradient purified synaptosomes were used for all glutamate release assays. In all cases, samples were diluted in HBM to a final concentration of 0.7 mg/ml, as determined by Protein-UV (Implen Nano Photometer). Suspension was spun down at 10000 g for 10 minutes (in tubes containing 1 ml) and pellets were kept on ice for a maximum of 4 hours. All steps were performed at 4°C. At no point during this procedure were synaptosomes subjected to hypertonic conditions. The pellet was thoroughly and carefully resuspended in warm HBM and incubated at 37°C for one hour before vitrification. During that time synaptosomes were treated with 0.1 _μ_M Calphostin C (CalC) for 30 min, 0.1 _μ_M Ro31-8220 for 30 min and 1 _μ_M Phorbol-12,13-dibutyrate (PDBu) for 5 min, as indicated in Figure S3. The lack of extracellular Ca^2+^in this preparation is in contrast to the resuspensions in HBM + 1.2mM CaCl_2_ that were used for synaptosomes from Munc13- and SNAP25-deficient slice cultures.

### Glutamate release assay

The glutamate release assay was performed according to (Nicholls et al., 1987) with modifications. Briefly, the synaptosomal pellet obtained using the Percoll gradient procedure was resuspended in Ca^2+^-free HBM +1 mg/ml BSA to a protein concentration of 0.7 mg/ml and incubated at 37 °C for 60 minutes.

1 mM NADP+, and 50 U/ml of glutamate dehydrogenase (GDH) were added 120 s before the measurements started. The fluorescence, resulting from reduction of NADP+ to NADPH that was caused by glutamate oxidation by GDH was measured in a spectrometer (Luminescence Spectrometer LS 50 B, Perkin Elmer, Waltham, USA) for 600 or 700 seconds. Excitation and emission wavelengths were 340 nm and 460 nm, respectively, and the sample was stirred in glass cuvettes. For calibration, 200 s before termination of the measurement, 2 _μ_M of glutamate was added as a reference. The calibration was done for each trace individually.

For measurements shown in Fig S3A, synaptosomes were treated with 0.1 _μ_M Calphostin C (CalC) for 30 min, 0.1 _μ_M Ro31-8220 (Ro31) for 30 min, 1 _μ_M Bisindolylmaleimide 1 (BIM-1) for 30 min and 1 _μ_M Phorbol-12,13-dibutyrate (PDBu) for 5 min, as indicated on the figure. At the end of the incubation period, synaptosomes were washed by 30 s centrifugation at 13000 rpm and resuspended in Ca^2+^-free HBM +1 mg/ml BSA. to a final protein concentration of 0.68 mg/ml. We added 1.33 mM CaCl_2_ at the beginning of the fluorescence measurements. Number of measurements for each condition was 5-7.

For measurements shown on Fig S3B, there were no pharmacological treatments performed during the incubation period. We added 0.7 mM CaCl_2_ 100 s after the beginning of the fluorescence measurement. In addition, 1 _μ_M PDBu was added at the beginning in some cases (labeled as PDBu) and was not added in other (labeled Control). In both cases, at 40 s after the onset of measurements, synaptosomes were treated with 5 mM KCl, 2.5-3 _μ_M Ionomycin, or they were not stimulated. Five measurements were done for each condition.

### RIM conditional knockout mice

Conditional RIM1/2flox/flox mice (RIM1flox:RIM2flox: RIMS1tm3Sud/J, RIMS2tm1.1Sud/J, Jackson Lab) (Kaeser et al., 2011) were used for all analyses. Mice were housed under a 12 h light-dark-cycle (light-cycle 7AM/7PM), in a temperature (22±2°C) and humidity (55±10%) controlled environment with food/water *ad libitum*. All procedures were planned and performed in accordance with the guidelines of the University of Bonn Medical Centre Animal-Care-Committee as well as the guidelines approved by the European Directive (2010/63/EU) on the protection of animals used for experimental purposes.

### Dissociated neuronal culture preparation from RIM conditional knockout mice

Mouse cortical neurons were prepared from RIM double floxed embryonic mice (E16-18) as previously described (Zürner et al., 2011). In brief, after dissection of cortices from embryonic mice and several rounds of washing with HBSS (Hank’s Balanced Salt Solution, Life Technologies), cells were digested with trypsin (0.025 g/ml, Life Technologies) for 20 min at 37°C. After several washing steps with HBSS, the remaining DNA was digested with DNase I (0.001 g/ml, Roche). Cannulas were used to dissociate the tissue and the suspension was passed through a Nylon cell strainer (100 _μ_m, BD Biosciences). Cells were seeded in a 10 cm dish coated with poly-D-lysine at a density of 1.2 million cells/dish or in a 6-well plate coated with poly-D-lysine at a density of 300,000 cells/well. Neurons were cultured in basal medium eagle (BME, Life Technologies) supplemented with 0.5% glucose (Sigma-Aldrich), 10% fetal calf serum (FCS, Life Technologies), 2% B-27 and 0.5 mM L-glutamine (Life Technologies) or Neurobasal medium (Thermo Fisher). Cells were maintained at 37°C in 5% CO_2_ until use. All embryos of a single mother were combined to maximize the number of neurons.

### Lentivirus production and infection of RIM double flox neuronal cultures

Lentiviruses were produced using a second-generation packaging system, as previously described (van Loo et al., 2019). In brief, 3 × 106 HEK293T cells (Clontech) were seeded on a 10 cm cell culture dish and transfected after 24 h with GenJet transfection reagent (SignaGen). Per dish 7.5 _μ_g packaging plasmid (psPax2, Addgene), 5 _μ_g VSV-G expressing envelope plasmid (pMD2.G, Addgene) and 4 _μ_g plasmid of interest (e.g. Cre-GFP) were used. After 12 h, transfection medium was replaced with DMEM containing Glutamax (Invitrogen) supplemented with 10% FBS. Transfected cells were incubated for 72 h to allow formation of viral vectors. Thereafter, the supernatant was filtered through 0.45 _μ_m PVDF membrane filters (GE Healthcare) to remove cell debris and other aggregates. To purify the virus, the filtered supernatant was layered on top of OptiPrepTM density gradient medium (Sigma-Aldrich) and centrifuged at 24,000 rpm for 2 h at 4°C using a SW-Ti32 swinging bucket (Beckman Coulter). The upper layer was discarded. The Opti-Prep-Layer, with the viral particles at its upper boundary, was mixed with TBS-5 buffer (containing in mM: 50 Tris-HCl, 130 NaCl, 10 KCl, 5 MgCl_2_). Viral particles were pelleted by centrifugation (24,000 rpm for 2 h at 4°C) and resuspended in TBS-5 buffer. Lentiviruses were stored at -80°C until use. Primary neurons cultured in 10 cm cell culture dishes were transduced with 7 _μ_l of lentiviral suspension per dish at DIV 2-6. For the transduction of primary neurons cultured in 6-well plates 3 _μ_l of lentiviral suspension were used per well. The primary cultured neurons were transduced with lentiviruses expressing active or inactive mutant-Cre recombinase to yield RIM double knock-out and RIM wild-type cells.

Lentiviral vectors encoding nuclear localization signal (NLS)–green fluorescent protein (GFP)–Cre and NLS–GFP-deltaCre were kindly provided by Thomas Südhof (Stanford University, Stanford, CA).

### Synaptosomal preparation from RIM conditional knockout cultures

RIM double floxed dissociated neuronal cultures (DIV 13 – 16), either treated with (GFP)–Cre or NLS– GFP-deltaCre expressing lentiviral particles at DIV2-6, were briefly rinsed with 2 ml of Tyrode’s buffer at room temperature, the buffer was removed, the dishes were placed on ice and 500 _μ_l of homogenization buffer (HB; 0.32 M sucrose and one tablet Complete mini EDTA-free protease inhibitor cocktail (Roche; 10 ml, pH 7.4)) (4°C) was added. The neurons were scraped off and placed in a glass homogenization tube (all at 4°C). Neurons were homogenized by one stroke at low speed (100rpm) and 7 strokes at 700rpm. The homogenate was centrifuged at 2000 g for 2 min (twice), the pooled supernatants were centrifuged for 12 min at 9500 g and the pellet (P2) was resuspended in Ca^2+^ free HBM (all at 4°C), yielding the crude synaptosomal fraction. The protein concentration was measured using a NanoDropTM spectrophotometer (ThermoFisher) and diluted with Ca^2+^ free HBM to little below 1 mg/ml. The sample was centrifuged at 10000g for 10min and the pellet was kept on ice until vitrification.

The above protocol was used for the data presented here. Despite several attempts to optimize the preparation, it has been extremely challenging to find synaptosomes that could be imaged by cryo-ET, for both RIM cDKO and RIM Ctrl. We also explored variations of this protocol, but they turned out to be less successful. Specifically, we used a filter (5_μ_m pore size) to remove large chunks (Chang et al., 2012) instead of homogenization and low speed centrifugation, purified the crude synaptosomal fraction on a 3/10/23% discontinuous Percoll gradient and varied the amount and the concentration of synaptosomes.

### Wild type dissociated neuronal culture

Primary neuronal culture was prepared according to the Banker method (Kaech and Banker, 2006) modified to grow cells on EM grids (Lucic et al., 2007). Gold Quantifoil grids (R1/4, Au 200 mesh, Quantifoil Micro Tools) were additionally coated with a 20-25 nm carbon layer (MED 020, BAL-TEC). For sterilization, grids were transferred to 4-well-dishes (Falcon, Amsterdam, Netherlands) and UV-irradiated for 30 minutes on a sterile bench. EM grids were coated with poly-L-lysine in a 1 mg/ml solution in 0.1 M borate buffer (pH 8.5) overnight in the dark as well as glass bottom dishes for light microscopy. Unbound poly-L-lysine was washed multiple times with autoclaved MilliQ water and the grids were soaked in neuronal plating medium until cell seeding.

To make an astroglial cell feeder culture, postnatal day 1 pups of Sprague-Dawley rats were sacrificed according to the guidelines of the Max-Planck-Institute of Biochemistry. After removing meninges, hippocampi and cortices were dissected and transferred to ice cold CMF-HBSS-Hepes (Calcium- and Magnesium-free Hank’s Balanced Salt Solution with 5% Hepes, Invitrogen, Carlsbad, USA). Tissue was minced and digested in 0.25% Trypsin and 0.1% (wt/vol) DNase I in a 37 °C water bath for 15 minutes and triturated with a 10-ml automated pipette. Cells were passed through a 70 _μ_m Cell strainer (Becton Dickinson and company, Heidelberg), centrifuged at 120 g for 10 minutes to remove enzymes and lysed cells and resuspended in Glial Medium (Minimal Essential Medium with Earl’s Salt and L-glutamine, 0.6% D-glucose, 100 U/ml Penicillin-Streptomycin, 10% fetal bovine serum; Invitrogen, Carlsbad, USA). Glial cells were plated in 75-cm^2^-flasks (Falcon, Amsterdam, Netherlands) at a density of 7.5 × 106 cells per flask and incubated in a CO_2_ incubator at 37 °C. Medium was exchanged every third day and flasks were swirled harshly to remove loosely attached cells from the flask surface such as microglia or O2A progenitor cells. Nearly confluent astroglia were harvested by trypsination, centrifuged at 120 g for 7 minutes and resuspended in Glial Medium for seeding in 6 mm dishes at a concentration of 1.0 × 105 cells per dish. Glial medium was exchanged every 3-4 days and replaced by Neurobasal/B27 medium for preconditioning. Preconditioned Neurobasal/B27 is required to feed primary cultured neurons.

Rat hippocampal neurons were prepared from Sprague-Dawley embryonic rats (E17-21) in accordance with the guidelines of the Max-Planck-Institute of Biochemistry. Hippocampi were dissected, transferred to ice cold CMF-HBSS-Hepes, minced, transferred to 0.25% Trypsin and 0.1% (wt/vol) DNase I in CMF-HBSS and incubated for digestion for 15 minutes at 37 °C. To wash out enzymes, CMF-HBSS medium was replaced three times and tissue was triturated. To remove left over clumps of biological material, the cell solution was passed through a 70 _μ_m cell strainer (Becton Dickinson and company, Heidelberg). After centrifugation (120 g, 10 min) cells were resuspended in neuronal plating medium (NPM: Minimal Essential Medium with Earl’s Salt and L-glutamine, 0.6% D-glucose, 5% fetal bovine serum; Invitrogen, Carlsbad, USA) and seeded to a concentration of 3.0 × 105 cells per well on EM grids and glass bottom dishes (MatTek corp., Ashland, USA) for immunohistochemistry. After 4 hours of settling cells on the substrate in a CO_2_ incubator, preconditioned Neurobasal/B-27 medium was added. Three days after cell seeding, 5 _μ_M AraC in preconditioned Neurobasal/B27 was added to each culture dish. One third of medium was exchanged once a week with Neurobasal/B27 and preconditioned Neurobasal/B27 in equal parts. At DIV 21, just before vitrification, cultures were treated by 1 _μ_M PDBu (final concentration) for 2 min.

### Vitrification of synaptosomes from Munc13- and SNAP25-deficient slice cultures and RIM conditional knockout cultures

Holey gold EM grids (Quantifoil R 2/1, 200 mesh; Quantifoil, Jena, Germany) were plasma cleaned in a Turbo Sputter Coater Med 010 (Balzers) for 2 min. To make fiducial marker solution, BSA-coated 10nm gold nanoparticle solution (Aurion, BSA tracer Conventional 10nm reagent, Product Code: 210.133, Wageningen, Netherlands) was concentrated by two centrifugations at 16000 g for 25 min (at 4°C) and the pellet was resuspended first in Ca^2+^-free HBM and the second time in HBM + 1.2mM CaCl_2_. Warm fiducial marker solution was mixed with synaptosomes at 1:10 ratio. 4 _μ_l of the mixture was deposited on each grid, allowed settle down for approximately 7 s, blotted at a small angle for 5-7 s with a filter paper (Whatman filter paper 1 Qualitative circles 90mm diameter, Cat No 1001 090) and vitrified by plunge freezing into liquid nitrogen (LN_2_)-cooled pure ethane (Westfalen AG, Ethane 2.5 99.5% Vol. % C2H6, Münster, Germany) using a portable manual plunger (designed and built by Max Planck’s workshop). Vitrified grids were stored in LN_2_ until EM imaging.

### Vitrification of neocortical synaptosomes

Holey copper or molybdenum EM grids (Quantifoil R 2/1 200 mesh; Quantifoil, Jena, Germany) were glow discharged (Harrick Plasma cleaner PDC-3XG) for 45 seconds. To make fiducial marker solution, BSA-coated 10nm gold nanoparticle solution (Aurion, BSA tracer Conventional 10nm reagent, Product Code: 210.133, Wageningen, Netherlands) was concentrated four times by two centrifugations at 14000g for 60 min and the pellet was resuspended in Ca^2+^-free HBM. Warm fiducial marker solution was mixed with synaptosomes at 1:10 ratio. 4 _μ_l of the mixture was deposited on each grid and vitrified by plunge-freezing into a liquid ethane/propane mixture using Vitrobot Mark III or Vitrobot Mark IV (Thermo Fisher Scientific). The vitrobot settings were: blot offset -3 (Mark III), blot force 10 (Mark IV), blotting time 10 s, at 37°C and 95% humidity. Vitrified grids were stored in LN_2_ until EM imaging.

### Vitrification of wild type neuronal cultures

Primary cultured neurons were seeded onto holey gold EM grids (Quantifoil R 1/4 gold 200 mesh). The fiducial marker solution was prepared as for neocortical synaptosomes. 4 _μ_l of the solution was applied to grids on which neurons were grown. They were vitrified using Vitrobot Mark IV (Thermo Fisher Scientific) using the same settings as for neocortical synaptosomes, except that a waiting time of 5 seconds was imposed to allow fiducial markers to diffuse on the sample. Vitrified grids were stored in LN_2_ until EM imaging.

### Cryo-FIB

Vitrified WT neuronal cultures grown on EM grids were thinned by focused ion beam (FIB) using Quanta 3D FEG and Scios DualBeam, FEI dual-beam microscopes, equipped with an 360° rotatable cryo-stage operated at -180 °C, as described before (Fukuda et al., 2014). Grids were mounted in FEI Autogrids (FEI), modified for shallow milling angles (Rigort et al., 2012), and placed in a cryo-FIB shuttle (Rigort et al., 2010). The additional carbon layer evaporated on the grids before plating neuronal cultures helped finding the correct orientation. The cryo-FIB shuttle was transferred to the microscope using a cryo-transfer system (PP3000Q, Quorum, East Sussex, UK). For initial experiments grids were sputtered (10 mA, 60 s) with a platinum layer to prevent surface charging of specimen. The effect of uneven milling (curtaining) was avoided by applying a protective layer of platinum onto the grid by a gas injection system (GIS). The region of interest was monitored by the electron beam at 5 keV and milled with 30 keV gallium ions. In general, rough milling (0.3 – 0.5 nA) was used to create rectangular holes surrounding the region of interest, and the milled area was further thinned (0.1 nA). Currents between 30 pA and 50 pA were used to polish milled surfaces. Alternatively, the cleaning cross section milling strategy was performed at 0.1 nA, where a selected area is milled line by line, using z-size of 6 _μ_m, 700 ns dwell time and 65% overlap.

Both the wedge and lamella FIB-milling strategies were employed. Wedges were milled in a relatively wide pattern of 35 _μ_m, typically at 5Q milling angle. Because of their shape, wedges contain only a small region close to the edge that is sufficiently thin for cryo-ET (Figure S6A). Also, the extended culturing time needed to ensure synaptogenesis often results in neurons growing processes on the bottom side of the EM grid carbon support (neurons are plated on the top side), thus making an additional thickness that cannot be removed by cryo-FIB milling because that would require unfeasibly low milling angles.

Lamellas, being more fragile than wedges, were milled to the width of 15 _μ_m. However, extended regions or even the entire length of a lamella can be milled to a suitable thickness. The milling was typically performed at 11Q and the desired lamella thickness just before the polishing step was 1 _μ_m. Because lamellas are more fragile than wedges, densely grown cultures are required for a sufficient support, but the increased sample thickness often caused unsatisfactory vitrification.

Furthermore, cryo-FIB thinning of neuronal processes, especially at low milling angles as required for wedges, resulted in material redeposition and surface contamination. Although we removed the surface contamination-induced 3D-reconstruction artefacts using a previously developed software (Fernandez et al., 2016), the contamination obscured the view to the interior of thinned regions, aggravating synapse detection.

### Cryo-ET acquisition and reconstruction

Tilt series were collected under a low dose acquisition scheme using SerialEM (Koster et al., 1997; Mastronarde, 2005) on Titan Krios and Polara microscopes (Thermo Fisher) equipped with a field emission gun operated at 300 kV, with a post-column energy filter (Gatan) operated in the zero-loss mode (20 eV slit width) and with a computerized cryostage designed to maintain the specimen temperature <-150°C. Tilt series were typically recorded from -60° to 60° with a 1.5° - 2° angular increment, using a modified version of the dose-symmetric scheme (Hagen et al., 2017) or in two halves starting from 0°, and the total dose was kept <100 e^-^/Å^2^. Almost all tilt series were recorded on a direct electron detector device (K2 Summit, Gatan) operated in the counting mode. Pixel sizes was 0.34 nm and 0.44 nm at the specimen level. Volta phase-plate with nominal defocus of 0.5-1 _μ_m (Danev et al., 2014) was used. Individual frames were aligned using Motioncor2 (Zheng et al., 2017). A few tilt series of neocortical synaptosomes were recorded on a 2kx2k charge-coupled device (CCD) camera (MegaScan, Gatan) at 5 _μ_m nominal defocus. Tilt series were aligned using gold beads as fiducial markers, and 3D reconstructions were obtained by weighted back projection (WBP) using Imod (Kremer et al., 1996). During reconstruction, the projections were binned using a factor of 4 (final pixel size 1.368 - 1.756 nm) and low pass filtered at the post-binning Nyquist frequency.

Cryo-FIB milling of neuronal cultures caused a strong redeposition of milled material. To remove 3D reconstruction artefacts induced by the redeposition, we applied the software procedure previously developed for this purpose (Fernandez et al., 2016). The number of iterations was set to five.

### Selection of tomograms

We selected for further processing tomograms that were of sufficient technical and biological quality. Specifically, tomograms were deemed technically acceptable if they did not contain any signs of ice crystal formation such as ice reflections or faceted membranes, and they had reasonable signal-to-noise ratio and proper tomographic alignment. We discarded synapses showing signs of deterioration such as elliptical small vesicles or strong endocytotic features. Only synaptosomes containing a presynaptic mitochondrion showing intact outer and inner membranes and cristae were kept. The selected synapses contained uninterrupted and smooth presynaptic and postsynaptic membranes, and the pre- and postsynaptic terminals were separated by a cleft of approximately 25 nm width, containing trans-cleft protein complexes.

### Detection, segmentation and analysis of tethers, connectors and SVs

SV-associated complexes, tethers and connectors, were detected and localized in an automated fashion and analyzed using Pyto package as described before (Fernández-Busnadiego et al., 2010; Lucic et al., 2016). In short, the AZ membrane was manually segmented in Amira (Visualization Sciences Group). SVs were segmented by manually tracing the maximum diameter profile and automatically extending it to a sphere.

Hierarchical connectivity segmentation was used to detect and localize connected clusters of pixels linking SVs to plasma membrane (tethers) and to other SVs (connectors). In this way, the segmented tethers and connectors contain the “core” structure, because they are segmented at the lowest greyscale density threshold at which they are detected.

For the analysis of vesicle distribution, the presynaptic cytoplasm (including SVs) was divided into 1-pixel-thick layers according to the distance to the AZ membrane, and the fraction of the layer volume occupied by SVs was measured. The surface concentration of SVs was calculated as the number of SVs in the proximal zone divided by the surface area of the AZ membrane. The SV distance to the AZ membrane was calculated as the shortest distance between the AZ membrane and SV pixels.

Analysis of detected tethers and connectors included the determination of their morphology, precise localization and their interrelationship. Connector and tether lengths were calculated as the minimal edge-to-edge distance between connector / tether voxels that contact an SV or plasma membrane, and they approximate the geodesic distance by taking into account central regions of tethers and connectors as previously described (Lucic et al., 2016). In this way curvature of tethers and connectors contributed to their calculated lengths. As expected, tether length was larger than the proximal SV distance to the AZ zone for all conditions (Figure S1B-E) because many tethers were curved, or they connected a side of a SV with the AZ membrane (Figure 3C). All features were analyzed within 250 nm from the AZ membrane.

To account for the missing wedge, we determined the synapse orientation as the direction of the vector perpendicular to the pre- and postsynaptic membranes in respect to the x-axis (these vectors were in x-y plane for all synapses). Even though there were no significant differences between the experimental conditions, we removed as many synapses as needed to equalize the mean angles between the conditions (synapses were removed in the order from the most extreme angle). This resulted in the removal of three SNAP25 KO, two Munc13 DKO +PDBu, two Plain (-Ca^2+^) and two Ro31-8220 synapses from all analysis steps that involved tethers and connectors.

All image processing and statistical analysis software procedures were written in Python and implemented in Pyto package (Lucic et al., 2016). Pyto uses NumPy and SciPy packages and graphs are plotted using Matplotlib (Harris et al., 2020; Virtanen et al., 2020; Hunter, 2007).

### Fitting atomic models

Tethers were extracted using Pyto package in two forms, directly from tomograms as greyscale densities (greyscale tethers), and as binary segments obtained by our hierarchical connectivity procedure (segmented tethers) (Lucic et al., 2016). Tethers were segmented in a fully automated manner, hence their isosurface levels were not manually adjusted for fitting.

Two atomic models were used for fitting: (i) the primed SNARE complex comprising one SNARE motif each of VAMP2 and Syntaxin 1a, two SNARE motifs of SNAP25, a helical fragment of Complexin 1 and the C2B domain of Synaptotagmin 1 [(Zhou et al., 2017) PDB id: 5w5d] (Figure 4A), and (ii) a Munc13 fragment, comprising C_1_, C_2_B and MUN domains that covers 55% of the entire Munc13 sequence [(Xu et al., 2017) PDB id: 5ue8] (Figure 4A).

We performed rigid body fitting of the selected atomic models, as follows. First, we performed a rough manual fit of atomic models in the segmented tethers, and then we used the automated fit using UCSF ChimearX software (Goddard et al., 2018). If needed, fits were adjusted manually. In all cases, we made sure that the global orientation of the atomic models was correct, namely, the C terminals of SNARE helices were localized close to the tether contacts with both SV and plasma membranes, and that N-terminal of the Munc13 C_1_C_2_BMUN model was oriented towards the plasma membrane and the C-terminal towards the SV.

Also, the atomic models were not allowed to overlap with the lipid membranes, examples of unsuccessful fits are shown in 4C. Finally, to avoid false negatives that might arise because tethers are segmented at the lowest greyscale level at which the tether is detected and thus may underestimate the real (greyscale) tether volume (Lucic et al., 2016), we determined whether the models could fit the greyscale tether.

In this way, we tested whether tethers were of sufficient size, and appropriate shape and orientation to accommodate the entire atomic models. A fit was considered satisfactory even if a part of the tether is not occupied by the model because it is possible that under native conditions other proteins bind those comprising the atomic models thus contributing to tethers, and because the Munc13 C_1_C_2_BMUN model contains 55% of the entire Munc13.

Specifically, we found that in all 25 cases where segmented short tethers were too small to accommodate SNARE complex helices (out of 80 short tethers), greyscale tethers provided an adequate fit. Fitting the Munc13 C_1_C_2_BMUN model into intermediate tethers of WT, Munc13 DHet, WT+PDBu and SNAP25 KO synapses, 30 out of 210 greyscale tethers could not accommodate the model (39 / 210 when considering only segmented tethers). Considering only SNAP25 KO synapses, 5 out of 41 greyscale tethers could not accommodate the model (6 / 41 when considering only segmented tethers). Furthermore, greyscale tethers could not accommodate the mode in 5 / 41 cases (6 / 41 segmented tethers) in SNAP25 KO and in 16 / 31 (17 / 31 segmented tethers) in intact neurons. Because the fairly large N-terminal region of Munc13 (540 amino acids) is not included in the Munc13 C_1_C_2_BMUN model, its conformation with respect to the C_1_C_2_BMUN region is not known, hence a successful fitting of the Munc13 C_1_C_2_BMUN model into a tether constitutes only a necessary condition for the tether to contain Munc13. Therefore, the number of tethers that cannot accommodate Munc13 may be higher than what we determined.

### Statistical analysis

Statistical analysis was performed between the experimental groups using only planned, orthogonal comparisons. Specifically, Munc13 DKO was compared to Munc13 DHet, SNAP25 KO to SNAP25 WT, WT +PDBu to WT, Munc13 DKO +PDBu to Munc13 DKO, SNAP25 KO +PDBu to SNAP25 KO, and al three -Ca^2+^treatements (PDBu, Calphostin C +PDBu and Ro31-8220 +PDBu) to Plain (-Ca^2+^). The numbers of synapses, vesicles, connectors, and tethers analyzed for each category are shown in Table S1. The intended number of synapses per experimental group was 5-10. The actual numbers varied because of the combination of the factors: (i) tomograms were acquired in batches of the same condition, (ii) tomogram reconstruction and selection did not immediately follow the acquisition and (iii) all tomograms that satisfied the selection criteria were included in the analysis. For the analysis of properties pertaining to individual SVs, connectors and tethers (such as the SV distance to the AZ membrane, tether length and fraction of tethers/connectors having a certain property), values within experimental groups were combined. Bars on the graphs show mean values and error bars the standard error of the mean (sem). In cases a fraction of SVs or tethers is shown, the error bars represent sem between synapse means. We used Student’s t test for statistical analysis of values that appeared to be normally distributed (e.g., vesicle diameter) and K-W test (nonparametric) for values deviating from the normal distribution (e.g., number of tethers and connectors per vesicle). For frequency data (e.g., fraction of connected and non-connected vesicles), χ^2^ test was used. In all cases, confidence levels were calculated using two-tailed tests. The confidence values were indicated in the graphs by a single asterisk for P < 0.05, a double asterisk for P < 0.01, and a triple asterisk for P < 0.001.

## Supplementary material

Table S1, Figures S1-S6 and Supplementary videos 1-3.

## Acknowledgements

We would like to thank Gabriela J. Greif for critical reading of the manuscript, S. Opitz for excellent technical assistance and Christoph Klatt for technical assistance.

## Funding

This work was supported by the DFG (LU 1819/2-1, SS 820/4-1), a HFSP RGP0020/2019 grant and by the Max Planck Society.

## Author Contributions

C.I., B.C., S.S., and V.L. designed research; C.P., U.L., C.I., J.B., C.C. and J.J.F. performed experiments; C.P., U.L., C.C. and V.L. analyzed data; J.S.-P., S.S., N.B., B.H.C. and V.L. supervised research, J.S.-P., S.S., N.B., W.B. and V.L. provided resources and acquired funding; V.L., C.I., C.P., U.L., B.C. and N.B. wrote the manuscript; all authors edited the manuscript.

## Declaration of interests

The authors declare no competing interests.

## Data availability

All data needed to evaluate the conclusions in the paper are present in the paper and the Supplementary Materials. Representative cryo-electron tomograms are deposited at Electron Microscopy Data Bank (EMDB). The Pyto software package used and further developed for this study is available at GitHub (https://github.com/vladanl/Pyto).

## Supplementary Materials for

**Table S1:** Number of analyzed proximal SVs, tethers and proximal connectors. In cases where synapses were removed to equalize the mean angular orientation, the number of synapses before / after the removal are specified. The numbers of tethers and connectors always include only the retained synapses. “Munc13 and SNAP25” denote organotypic slice culture synaptosomes, “Pharmacologically treated” denote neocortical synaptosomes,”RIM”denote dissociated culture synaptosomes and”Neuronal culture”denote synapses from intact neurons.

|  | N synapses | N proximal SVs | N tethers | N proximal connectors |
| --- | --- | --- | --- | --- |
| Munc13 and SNAP25 |  |  |  |  |
| WT | 14 | 182 | 555 | 630 |
| WT +PDBu | 8 | 59 | 200 | 102 |
| Munc13 DHet | 8 | 80 | 266 | 253 |
| Munc13 DKO | 13 | 96 | 117 | 356 |
| Munc13 DKO +PDBu | 9 / 7 | 62 | 61 | 184 |
| SNAP25 WT | 9 | 110 | 349 | 340 |
| SNAP25 KO | 14 / 11 | 147 | 300 | 471 |
| SNAP25 KO +PDBu | 7 | 124 | 457 | 625 |
| Pharmacologically treated |  |  |  |  |
| Plain (-Ca<sup>2+</sup>) | 17 / 15 | 228 | 254 | 244 |
| PDBu (-Ca<sup>2+</sup>) | 16 | 164 | 270 | 164 |
| Calphostin C + PDBu (-Ca<sup>2+</sup>) | 9 | 87 | 107 | 101 |
| Ro31-8220 + PDBu (-Ca<sup>2+</sup>) | 8 / 6 | 118 | 167 | 121 |
| RIM |  |  |  |  |
| RIM Ctrl | 4 | 34 | 188 |  |
| RIM cDKO | 5 | 10 | 39 |  |
| Neuronal cultures |  |  |  |  |
|  | 5 | 39 | 108 |  |

Supplementary video 1: Cryo-ET image of a WT synapse, followed by 3D rendering of SVs (blue), the AZ membrane (blue), segmented tethers (violet) and connectors (green).

Supplementary video 2: The same as Supplementary video 1, except that a Munc13 DKO synapse is shown.

Supplementary video 3: The same as Supplementary video 1, except that a synapse from intact neuronal cultures is shown.

**Figure S1:**
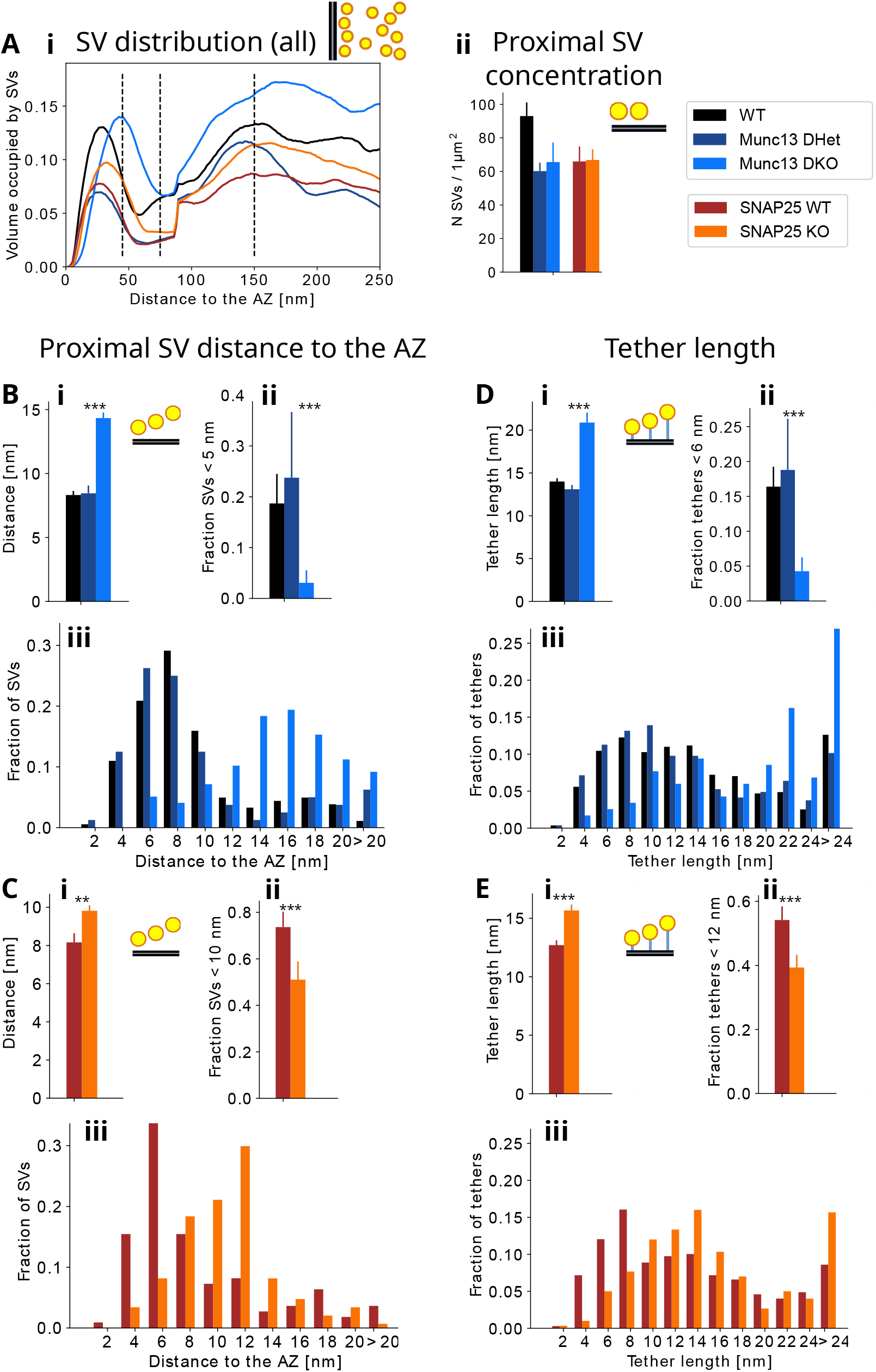
Influence of Munc13 and SNAP25 on the proximal SV location and tether length, additional data. (A) Mean SV distribution up to 250 nm to the AZ membrane (left) and the surface concentration of the proximal SVs at the AZ (right). (B) Proximal SV distance to the AZ membrane (above, left), fraction of proximal SVs located <5 nm to the AZ membrane (above, right) and the histogram of the SV distances (below), all from Munc13-related conditions. (C) Proximal SV distance to the AZ membrane (above, left), fraction of proximal SVs located <10 nm to the AZ membrane (above, right) and the histogram of the SV distances (below), all from SNAP25-related conditions. (D) Tether length (above left), fraction of tethers shorter than 6 nm (above right) and histogram of tether lengths (below), all from Munc13 conditions. (E) Tether length (above left), fraction of tethers shorter than 12 nm (above right) and histogram of tether lengths (below), all from SNAP25-related conditions.

**Figure S2:**
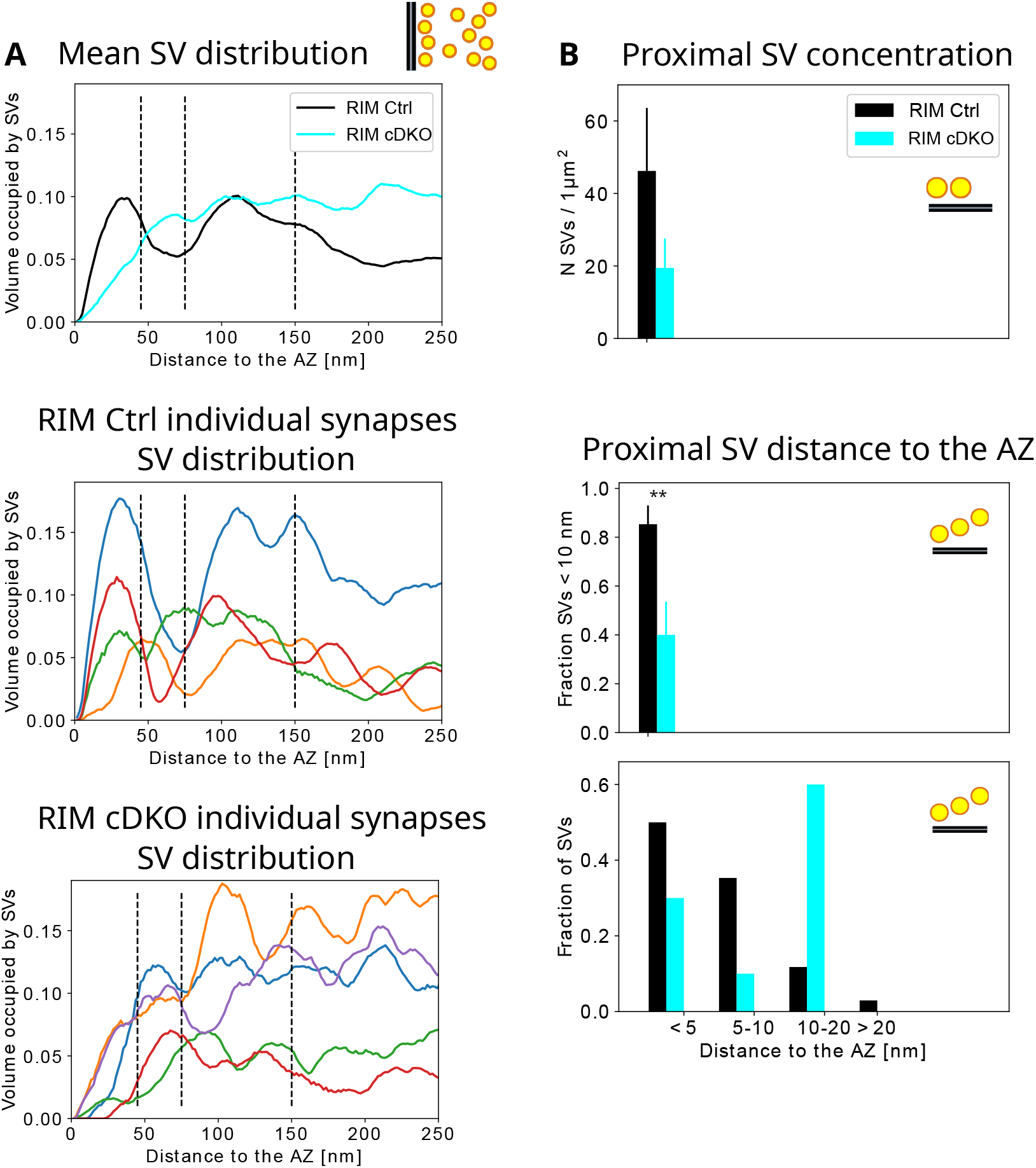
Role of the RIM family for SV localization. (A) SV distribution up to 250 nm to the AZ membrane, mean values (top), the individual WT synapses (middle) and the individual RIM cDKO synapses (bottom). Dashed black lines show separation between proximal intermediate and distal SVs. (B) Proximal SVs, the surface concentration at the AZ (top), fraction of SVs <10 nm to the AZ membrane (middle) and the histogram of the distances (bottom).

**Figure S3:**
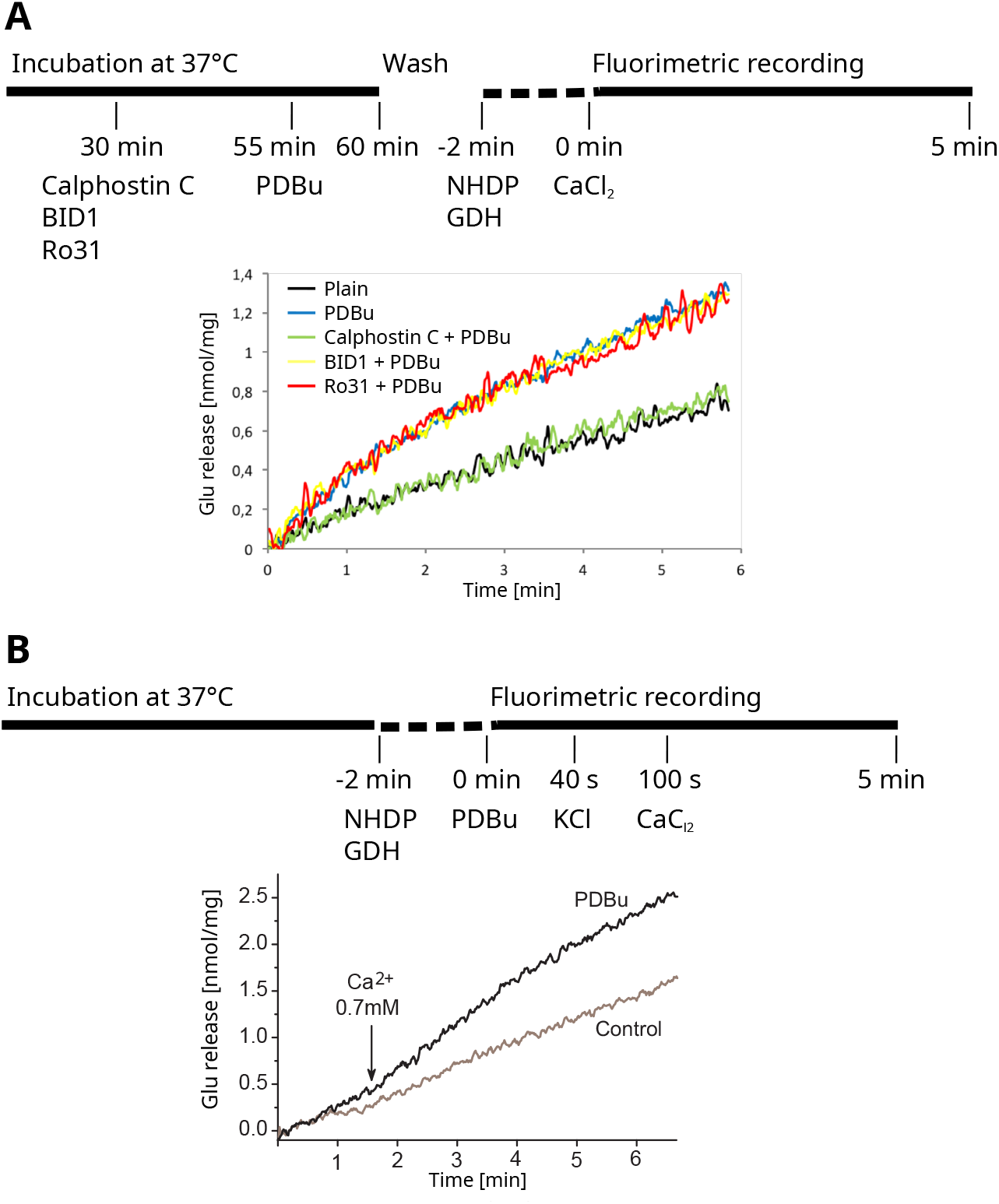
Glutamate release in neocortical synaptosomes. (A) Spontaneous release in pharmacologically treated synaptosomes (number of experiments 5-7 per treatment). (B) The influence of extracellular Ca^2+^ under a mild stimulation (5 mM KCl) and in the presence of PDBu. The exact timings are shown above the graphs.

**Figure S4:**
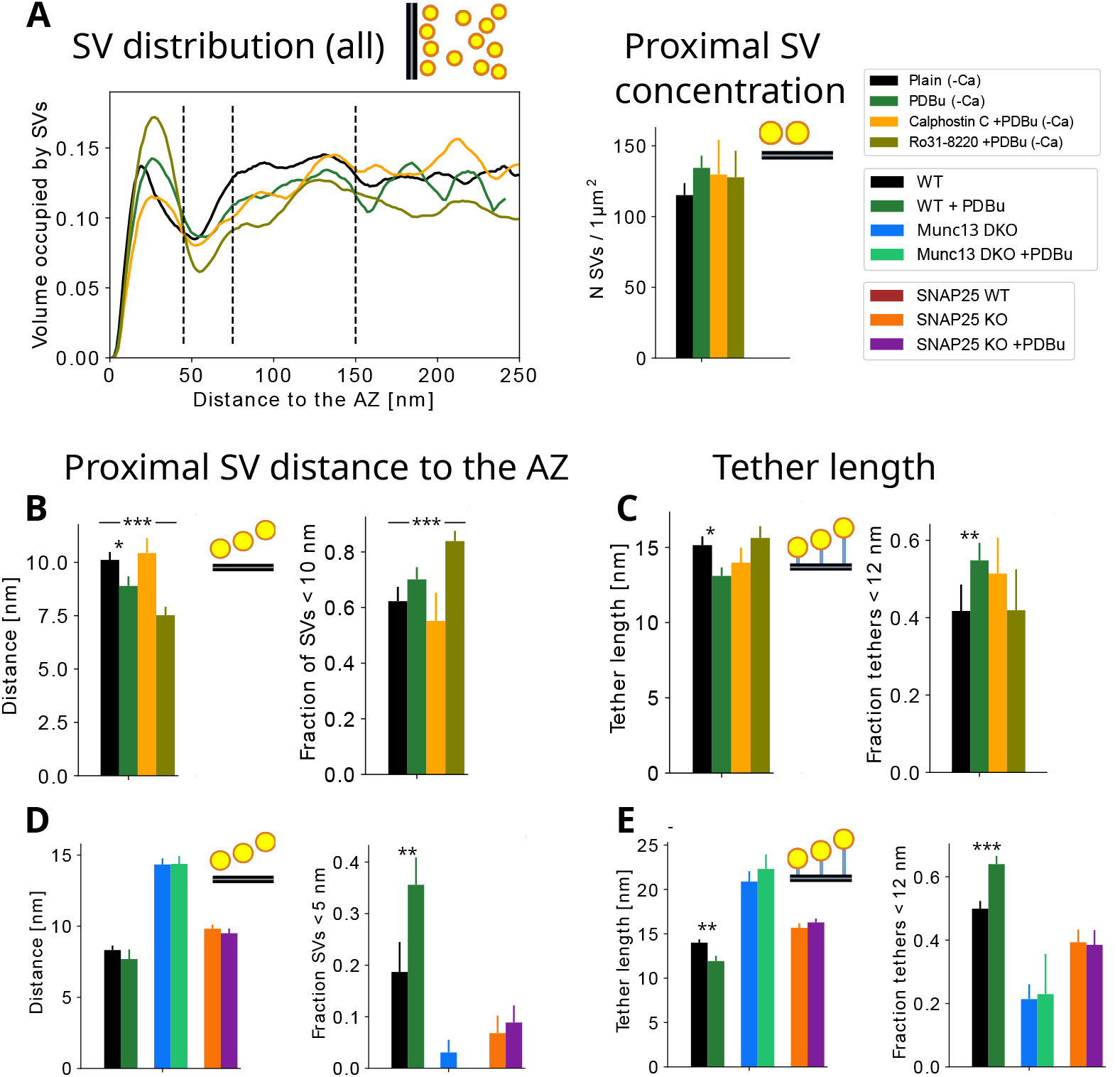
Influence of PDBu on SV distribution and tether length. (A) Mean SV distribution up to 250 nm to the AZ membrane (left) and the surface concentration of the proximal SVs at the AZ (right). (B) Proximal SV distance to the AZ membrane (left) and fraction of proximal SVs located <10 nm to the AZ membrane (right). (C) Tether length (left) and the fraction of tethers shorter than 12 nm (right). (D) Proximal SV distance to the AZ membrane (left) and fraction of proximal SVs located <5 nm to the AZ membrane (right). (E) Tether length (left) and the fraction of tethers shorter than 12 nm (right). (A-C) pharmacologically treated neocortical synaptosomes, (D-E) Munc13 and SNAP25 conditions.

**Figure S5:**
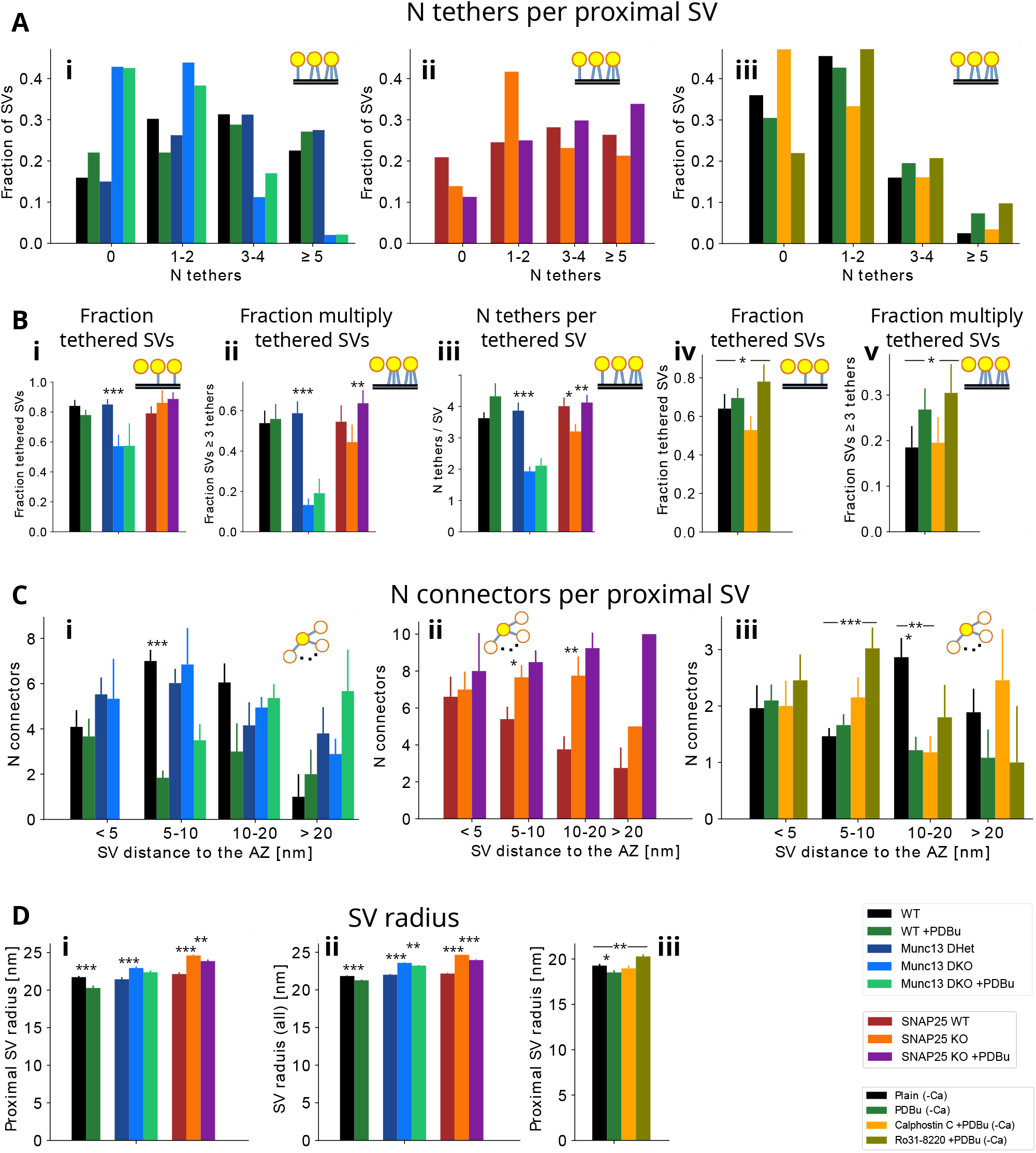
SV tethering, connectivity and size. (A) Histograms of the number of tethers per proximal SV. (B) Characterization of tethering, as indicated on the graphs. (C) The number of connectors per proximal SV for different SV classes. (D) SV radius for proximal SVs (left and right) and for all SVs within 250 nm to the AZ membranes (center). In all cases, the data for Munc13 and SNAP25 conditions are shown on the left and for the pharmacologically treated neocortical synaptosomes on the right.

**Figure S6:**
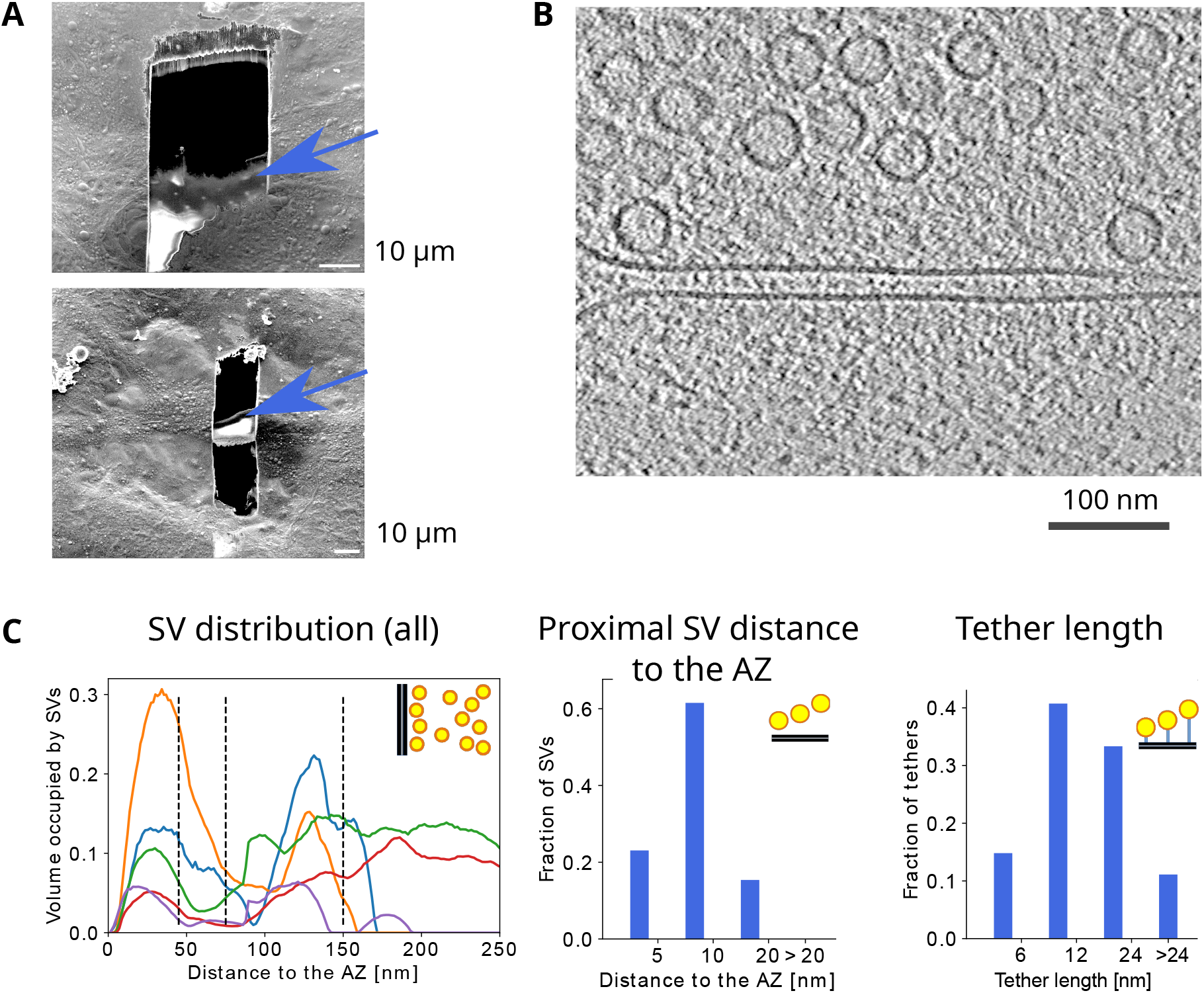
Intact neurons. (A) SEM images of a cryo-FIB milled wedge (above) and lamella (below). Arrows indicate the areas that are sufficiently thin for cryo-ET imaging. (B) Cryo-ET slice of a synapse. (C) SV distribution up to 250 nm to the AZ membrane of individual synapses (left). Histograms of the SV distances to the AZ membrane (center) and tether lengths (right).

## Notes

### Competing Interest Statement

The authors have declared no competing interest.

### Summary of Updates

This version of the manuscript has been revised to improve the presentation of the data.

